# Proteomic Analysis Reveals a PLK1-Dependent G2/M Degradation Program and Links PKA-AKAP2 to Cell Cycle Control

**DOI:** 10.1101/2023.10.11.561963

**Authors:** Ryan D Mouery, Carolyn Hsu, Thomas Bonacci, Derek L Bolhuis, Xianxi Wang, Christine A Mills, E Drew Toomer, Owen G Canterbury, Kevin C Robertson, Timothy B Branigan, Nicholas G Brown, Laura E Herring, Michael J Emanuele

## Abstract

Targeted protein degradation by the ubiquitin-proteasome system is an essential mechanism regulating cellular division. The kinase PLK1 coordinates protein degradation at the G2/M phase of the cell cycle by promoting the binding of substrates to the E3 ubiquitin ligase SCF^βTrCP^. However, the magnitude to which PLK1 shapes the mitotic proteome has not been characterized. Combining deep, quantitative proteomics with pharmacologic PLK1 inhibition (PLK1i), we identified more than 200 proteins whose abundances were increased by PLK1i at G2/M. We validate many new PLK1-regulated proteins, including several substrates of the cell cycle E3 SCF^Cyclin^ ^F^, demonstrating that PLK1 promotes proteolysis through at least two distinct SCF-family E3 ligases. Further, we found that the protein kinase A anchoring protein AKAP2 is cell cycle regulated and that its mitotic degradation is dependent on the PLK1/βTrCP-signaling axis. Interactome analysis revealed that the strongest interactors of AKAP2 function in signaling networks regulating proliferation, including MAPK, AKT, and Hippo. Altogether, our data demonstrate that PLK1 coordinates a widespread program of protein breakdown at G2/M. We propose that dynamic proteolytic changes mediated by PLK1 integrate proliferative signals with the core cell cycle machinery during cell division. This has potential implications in malignancies where PLK1 is aberrantly regulated.

## Introduction

Cell cycle progression enables duplication and segregation of the genome into identical daughter cells. A fundamental feature of the eukaryotic cell cycle is that it proceeds in a unidirectional and irreversible manner. The timely degradation of cell cycle regulatory proteins by the ubiquitin-proteasome system ensures cell cycle irreversibility, prevents genome reduplication, and maintains genome integrity (Dang et al. 2021). The ubiquitin system regulates protein degradation by utilizing a cascade of enzymes termed E1, E2, and E3 which culminates in the post-translational attachment of the small protein ubiquitin to target proteins. The formation of polyubiquitin chains on substrates is often a proteolytic signal that targets substrates to the proteasome, thereby triggering their degradation. E3 ubiquitin ligases are the enzymes responsible for directly binding to substrates and thus confer substrate specificity in the ubiquitin pathway (Hershko and Ciechanover 1998). Despite the importance of ubiquitin signaling in cell cycle regulation, very little is known about its role in G2/M phase. This is despite the importance of G2/M in proliferative decision making and cell cycle arrest in response to genotoxic stress (Spencer et al. 2013; Naetar et al. 2014; Min et al. 2020; Yang et al. 2017).

The Skp1-Cul1-F box (SCF) family of E3 ubiquitin ligases plays critical roles in cell cycle progression (Jang et al. 2020; Cardozo and Pagano 2004). The SCF relies on a family of ∼70 substrate receptors termed F-box proteins which recruit substrates to the SCF for ubiquitination (Nguyen and Busino 2020; Bai et al. 1996). The F-box proteins βTrCP and Cyclin F, among others, play critical roles in cell cycle progression (Zheng et al. 2016). βTrCP1 and βTrCP2 (hereafter βTrCP, unless otherwise noted) are functionally redundant and remain constitutively active throughout the cell cycle to control the degradation of many different proteins (Paul et al. 2022). βTrCP recognizes substrates containing a consensus DSGxx(x)S phospho-degron (Winston et al. 1999) following phosphorylation by upstream kinases (Hunter 2007). Cyclin F is the founding member of the F-box family and its expression is cell cycle regulated (Bai et al. 1994). While it contains a cyclin homology domain that binds to Cy-motifs in substrates, Cyclin F neither binds nor activates a cyclin dependent kinase. Instead, Cyclin F drives cell cycle progression via ubiquitination (D’Angiolella et al. 2013). Both Cyclin F and βTrCP control the destruction of many cell cycle proteins, and thus, play key roles in proliferation.

Polo-like kinase 1 (PLK1) is a multi-functional cell cycle kinase which is most highly active during G2-phase and mitosis. The activation of PLK1 at G2 helps promote mitotic entry in many cell types through the coordinated phosphorylation of myriad substrates (van Vugt et al. 2004; Macůrek et al. 2008; Bruinsma et al. 2012; Gheghiani et al. 2017). Once cells enter mitosis, PLK1 plays vital roles in controlling spindle organization, microtubule dynamics, and kinetochore function. Later in mitosis, PLK1 promotes mitotic exit and cytokinesis, altogether highlighting PLK1’s critical role in cell division (Petronczki et al. 2008). PLK1 has also been shown to coordinate protein degradation at mitotic entry. Established PLK1-dependent degradation substrates include EMI1 (Hansen et al. 2004; Moshe et al. 2004) and USP37 (Burrows et al. 2012), both of which antagonize the cell cycle E3 APC/C; WEE1 (Watanabe et al. 2004), which controls cyclin dependent kinase activity; and Claspin (Peschiaroli et al. 2006; Mailand et al. 2006; Mamely et al. 2006), which promotes DNA damage checkpoint signaling in S-phase. Additionally, PLK1-mediated degradation of Bora promotes normal progression through mitosis by regulating the function of Aurora Kinase A (Seki et al. 2008; Chan et al. 2008). PLK1 regulates the degradation of these proteins through the phosphorylation of the aforementioned degron sequence, allowing for binding to SCF^βTrCP^. Since PLK1 activity is cell cycle regulated (Golsteyn et al. 1995; Akopyan et al. 2014), there is a limited temporal window in which the degradation of these substrates is controlled.

Despite our knowledge of the PLK1/βTrCP-signaling axis and the importance of PLK1 in normal and cancer cell cycles, the magnitude to which PLK1 shapes the mitotic proteome through regulated protein degradation has not been studied. Determining the extent to which PLK1 controls proteome dynamics will shed light on mechanisms of G2/M control and the potential clinical use of PLK1 inhibitors.

In this study, we used mass spectrometry to perform global, quantitative proteomic analysis on mitotically synchronized, PLK1-inhibited colorectal cancer cells that harbor activating mutations in the oncogenic small GTPase KRAS. We identified over 200 proteins whose abundances are negatively regulated by PLK1 activity. Together with the validation of many new substrates, this demonstrates a widespread, PLK1-regulated G2/M degradation program. Significantly, we found that the degradation of most known SCF^Cyclin^ ^F^ substrates is PLK1-dependent, suggesting that PLK1 coordinates degradation through at least two cell cycle E3 ubiquitin ligases.

Among the many proteins validated was A-kinase anchor protein 2 (AKAP2), which we show to be degraded in mitosis in a PLK1- and βTrCP-dependent manner. AKAP family proteins bind to the regulatory subunit of protein kinase A (PKA) and function as scaffolds to facilitate localized PKA signaling within distinct areas of the cell (Omar and Scott 2020). There are over 50 members of the AKAP family and these are implicated in diverse aspects of cell physiology (Han et al. 2015). However, the functions of AKAP2 are poorly understood and it has mainly been studied in specialized tissues such as chondrocytes (Wang et al. 2021), cardiomyocytes (Maric et al. 2021), ocular lens (Gold et al. 2012), and various cancer types (Li et al. 2017; Thakkar et al. 2016). AKAPs have not been shown to undergo regulated proteolysis during cell cycle progression. Interestingly, mass-spectrometry (MS) analysis revealed that AKAP2 interacts with several proteins involved in regulation of the cell cycle as well as proteins functioning within signaling networks regulating cellular proliferation, such as the MAPK, AKT, and Hippo pathways.

Altogether, our data highlight new roles of protein degradation in the G2 phase of the cell cycle and provide new connections between PLK1, PKA, and various signaling pathways regulating proliferation. We previously showed that PLK1 inhibition or depletion has a synthetic lethal relationship with activating mutations in KRAS in colorectal cancer cells (Luo et al. 2009). PLK1 inhibitors are under clinical investigation and are being specifically tested in KRAS mutant colorectal cancers (Chiappa et al. 2022; Iliaki et al. 2021). Thus, our findings may have important implications for the treatment of cancer.

## Results

### PLK1 Regulates a G2/M Degradation Program

In order to understand how the mitotic proteome is affected by PLK1 activity, we arrested HCT116 cells in prometaphase using nocodazole in the presence of DMSO or either of two distinct small molecule inhibitors targeting PLK1 (BI2536 and BI6727). After treatment, synchronized cells were collected using mitotic “shake-off” and whole-cell extracts were prepared for label-free, quantitative proteomics using data independent acquisition (**Figure 1A** **and Table S1)**. In parallel, we analyzed asynchronous HCT116 cells and compared these to the DMSO-treated mitotically arrested cells. Validating our synchronization, we identified many key mitotic proteins among those most strongly upregulated in mitosis. These include Cyclin B1, Aurora Kinase A, components of the Chromosomal Passenger Complex (AURKB, BIRC5/Survivin, INCENP and CDCA8/Borealin), the centromere/kinetochore proteins CENP-F, BUB1, Shugoshin 1 and Shugoshin 2, and PLK1 (**Figure 1B** **and Table S1)**.

**Figure 1.**
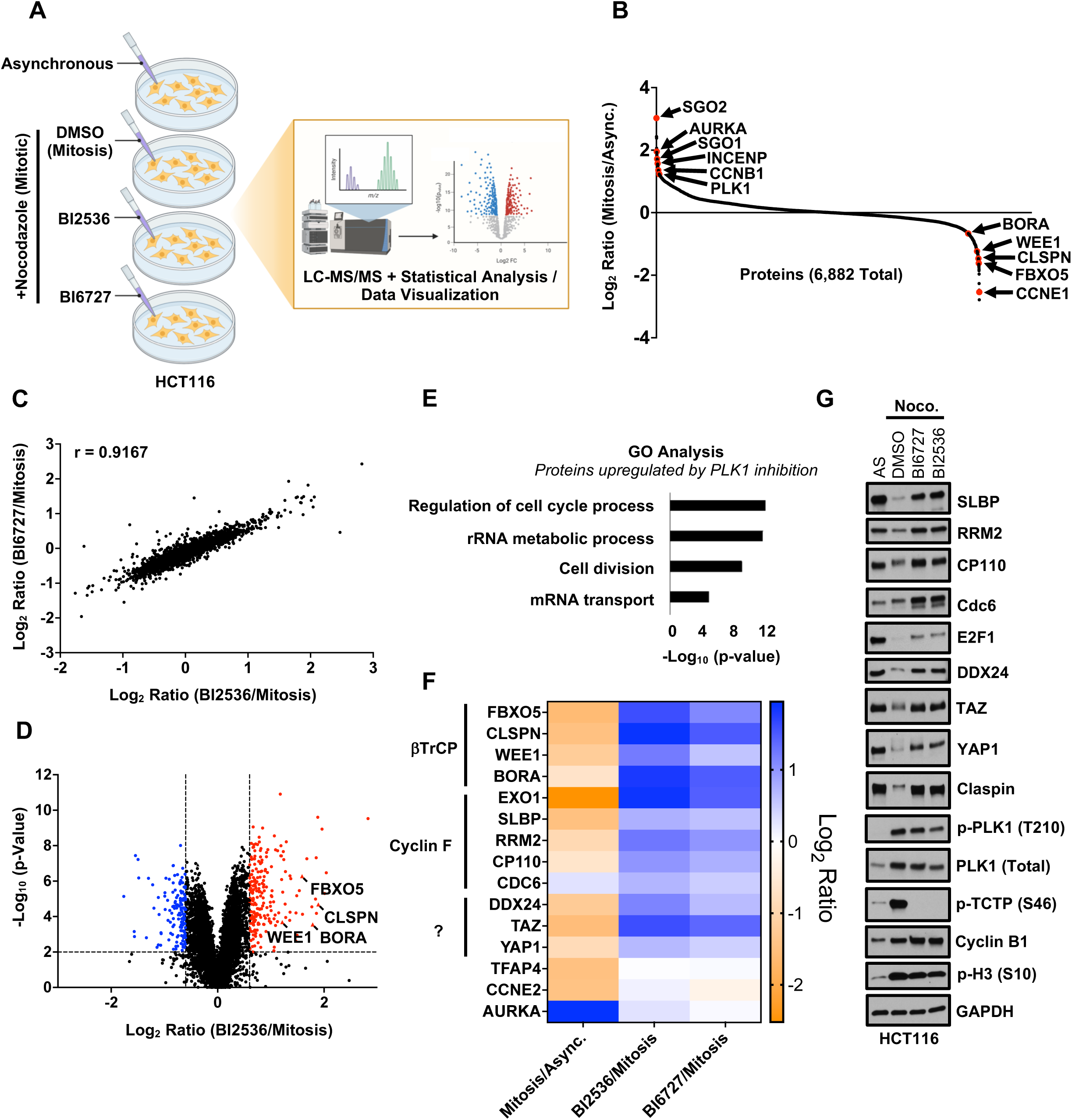
PLK1 Regulates a G2/M Degradation Program. (A) Schematic describing the workflow for the PLK1i proteomics experiment. HCT116 cells were grown asynchronously or were co-treated with 100 ng/mL nocodazole plus DMSO, 100 nM BI2536, or 100 nM BI6727 for 16 hours. After 16 hours, mitotic-arrested cells were collected using mitotic “shake-off” procedure. All samples were then prepared for label free, quantitative LC-MS/MS analysis using data independent acquisition (DIA). (B) Plot showing the Log2 ratio of all proteins identified by mass-spectrometry when comparing samples from DMSO-treated mitotic cells to samples from asynchronously growing cells. Each dot along the x-axis represents an individual protein. Proteins known to be up-regulated or down-regulated in mitosis are indicated. (C) Plot showing the Log2 ratio of all proteins identified by mass-spectrometry when comparing samples from BI2536-treated mitotic cells to samples from DMSO-treated mitotic cells (x-axis) vs. the Log2 ratio of all proteins identified by mass-spectrometry when comparing samples from BI6727-treated mitotic cells to samples from DMSO-treated mitotic cells (y-axis). (D) Volcano plot showing the Log2 ratio (x-axis) and –Log10 (p-value) (y-axis) of all proteins identified by mass-spectrometry when comparing samples from BI2536-treated mitotic cells to samples from DMSO-treated mitotic cells. Significant fold-change cutoffs were set at Log2 = -0.6 or Log2 = 0.6 and statistical significance was set at –Log10 (p-value) = 2 (p = 0.01) which are denoted by the dashed lines. Positive controls are labeled and indicated as triangle dots. (E) GO analysis highlighting top biological processes enriched among proteins upregulated in mitosis following PLK1 inhibition. (F) Heatmap showing the Log2 ratio of indicated proteins (left) identified by mass-spectrometry when comparing samples from DMSO-treated mitotic cells to samples from asynchronously growing cells (column 1), when comparing samples from BI2536-treated mitotic cells to samples from DMSO-treated mitotic cells (column 2), or when comparing samples from BI6727-treated mitotic cells to samples from DMSO-treated mitotic cells (column 3). Known substrates of βTrCP or Cyclin F are clustered together. The E3 ubiquitin ligases for DDX24, WWTR1 (TAZ), and YAP1 during mitosis are unknown and are also clustered together. TFAP4 is a known βTrCP substrate that is not dependent on PLK1 for degradation. CCNE2 and AURKA are cell cycle regulated proteins whose expression is not affected by PLK1 inhibition. (G) Validation of mass-spectrometry data. HCT116 cells were treated and collected as in (A) and samples were prepared for and analyzed by immunoblot for the indicated proteins.

To understand how PLK1 impacts total protein levels in mitosis, we focused on changes in mitotic cells following PLK1 inhibition (PLK1i). Data from the two PLK1 inhibitors were strongly correlated (Pearson’s correlation coefficient = 0.9167), suggesting that the changes seen are not simply off-target effects of the small molecules (**Figure 1C**). Using cutoffs of 0.6 for Log_2_ Ratio (PLK1i/Mitosis) and a p-value of 0.01 for statistical significance, we identified more than 200 proteins whose abundances were upregulated in PLK1-inhibited mitotic cells compared to control mitotic cells (**Figure 1D** **and Figure S3B)**. Among the proteins upregulated in mitosis in response to PLK1i, we identified WEE1, Claspin, Bora, and FBXO5/EMI1, whose mitotic degradation has been previously shown to be PLK1-dependent. We performed gene ontology (GO) analysis on the proteins upregulated in response to PLK1i and found that “regulation of cell cycle process” and “cell division” were among the most enriched GO-terms, consistent with PLK1’s role as a cell cycle regulator (**Figure 1E** **and Table S2)**.

Established PLK1/βTrCP substrates were strongly downregulated in mitosis when compared to asynchronous cells and strongly upregulated in PLK1-inhibited mitotic cells compared to control mitotic cells (**Figure 1F**). We therefore searched our dataset for additional proteins following this pattern, as these are strong candidates for potentially new PLK1-dependent proteasomal targets at G2/M phase. This analysis identified the RNA helicase DDX24 and transcriptional regulators YAP1 and TAZ as downregulated in mitosis in a PLK1-dependent manner, and all three were validated by endogenous immunoblot (**Figure 1G**). Interestingly, this analysis also identified many known substrates of the cell cycle E3 ubiquitin ligase SCF^Cyclin^ ^F^ (**Figure 1F**). In HCT116 cells, we validated by immunoblot of endogenous proteins that the mitotic regulation of the Cyclin F substrates SLBP, RRM2, CP110, Cdc6, and E2F1 is PLK1-dependent (**Figure 1G**). Importantly, PLK1 inhibition did not result in non-specific changes to cell cycle proteins. Phosphorylation of TCTP on S46 serves as a marker of PLK1 activity (**Figure 1G**). Cell synchrony was confirmed by immunoblotting for total cyclin B, phosphorylation of PLK1 within its activation loop (residue T210), and phosphorylation of Histone H3 on S10 (**Figure 1G**). Together, these data indicate that the differences in protein abundances seen are a result of PLK1 inhibition and not a result of unequal synchronization in mitosis.

To test whether the regulation of these protein abundances occurs in multiple cell types, we next examined HeLa cells arrested in mitosis with or without PLK1i. We observed a similar pattern of PLK1 dependence for the regulation of Cyclin F substrates, suggesting that this regulation is not cell line-dependent **(Figure S1A)**. We also demonstrated that the regulation of Cyclin F substrates by PLK1 is indeed Cyclin F-dependent, and not through βTrCP or another E3. The abundance of Cyclin F substrates in mitosis was unchanged in Cyclin F knockout cells following PLK1 inhibition **(Figure S1B/C)**, suggesting an epistatic relationship in which PLK1 functions upstream of Cyclin F to promote the degradation of its substrates.

Interestingly, unlike in the case of βTrCP, Cyclin F does not directly recognize phospho-degrons in substrates (Enrico et al., 2021). This raises the question as to how PLK1 promotes the degradation of Cyclin F substrates. We hypothesized that PLK1 may regulate Cyclin F itself, rather than its individual substrates. Consistently, we found that transient PLK1 inhibition resulted in the loss of Cyclin F abundance across six different cell lines **(Figure S2A)**. Further, the abundance of exogenously expressed Cyclin F increased when co-expressed with PLK1 **(Figure S2B)**, suggesting that PLK1 positively regulates Cyclin F abundance. To gain insight into how PLK1 regulates Cyclin F abundance, we relied on PLK1 inhibition **(Figure S2C/D)** or PLK1 over-expression **(Figure S2E/F)** and performed cycloheximide chase analysis. PLK1i resulted in decreased Cyclin F stability whereas PLK1 over-expression resulted in a marked increase of Cyclin F stability, suggesting that PLK1 regulates Cyclin F through a post-translational mechanism.

Altogether, these data suggest that PLK1 regulates a G2/M degradation program in coordination with at least two distinct E3 ubiquitin ligases.

### AKAP2 is a Cell Cycle Regulated Protein

Most proteins whose degradation are controlled by PLK1 are cell cycle regulated and their expression is decreased in mitosis. Among the most significantly downregulated proteins in mitosis was A-kinase anchor protein 2 (AKAP2) which demonstrated a 2.6-fold decrease in mitosis (**Figure 2A**). AKAP2 also showed a 3.9-fold or 3.5-fold increase in mitotic cells treated with BI2536 or BI6727, respectively, when compared to control mitotic cells **(Figure S3A/B)**. Although AKAP2 has not been described previously to be regulated in a cell cycle dependent manner, we found that AKAP2 protein level dynamics were remarkably similar to Claspin, whose phosphorylation by PLK1 in G2/M triggers its ubiquitination and degradation (**Figure 2B**). We therefore hypothesized that AKAP2 is a cell cycle regulated protein whose mitotic degradation is regulated by PLK1.

**Figure 2.**
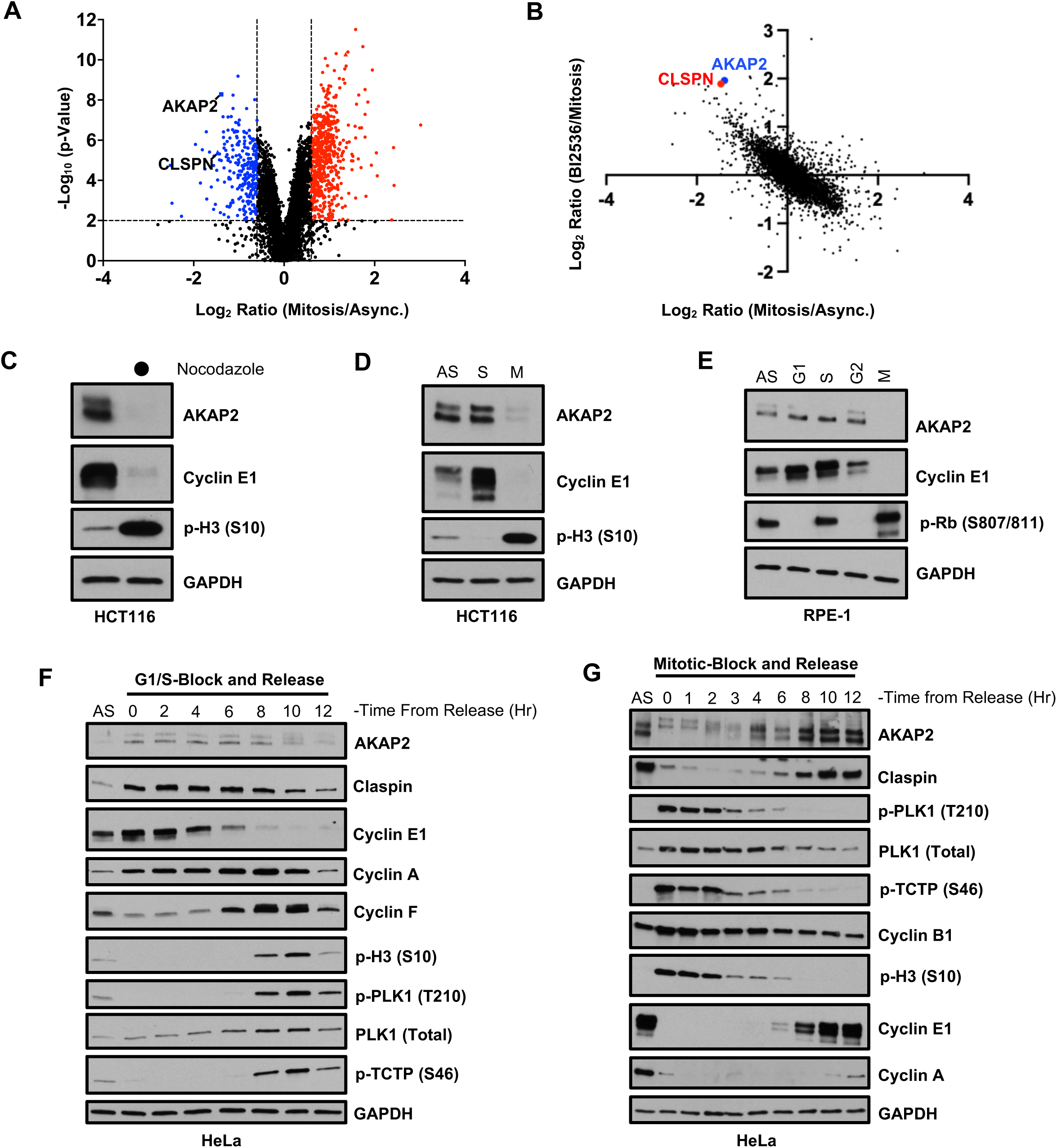
AKAP2 is a Cell Cycle Regulated Protein. (A) Volcano plot showing the Log2 ratio (x-axis) and –Log10 (p-value) (y-axis) of all proteins identified by mass-spectrometry when comparing samples from DMSO-treated mitotic cells to samples from asynchronously growing cells. Significant fold-change cutoffs were set at Log2 = -0.6 or Log2 = 0.6 and statistical significance was set at –Log10 (p-value) = 2 (p = 0.01) which are denoted by dashed lines. Claspin is identified as a positive control. (B) Plot showing the Log2 ratio of all proteins identified by mass-spectrometry when comparing samples from DMSO-treated mitotic cells to samples from asynchronously growing cells (x-axis) and the Log2 ratio of all proteins identified by mass-spectrometry when comparing samples from BI2536-treated mitotic cells to samples from DMSO-treated mitotic cells (y-axis). Claspin is identified as a positive control. (C) HCT116 cells were grown asynchronously or synchronized in mitosis by treatment with 100 ng/mL nocodazole for 16 hours. Mitotic-arrested cells were collected using mitotic “shake-off” procedure. All cell lysates were analyzed by immunoblot for the indicated proteins. (D) HCT116 cells were grown asynchronously, synchronized in S-phase (2 mM thymidine) or synchronized in mitosis (100 ng/mL nocodazole) for 16 hours. Mitotic-arrested cells were collected using mitotic “shake-off” procedure. All cell lysates were analyzed by immunoblot for the indicated proteins. (E) RPE-1 cells were grown asynchronously or synchronized in G1 (1 μM Palbociclib), S (2 mM Thymidine), G2 (10 μM RO-3306) or mitosis (100 ng/mL nocodazole) for 20 hours. Mitotic-arrested cells were collected using mitotic “shake-off” procedure. All cell lysates were analyzed by immunoblot for the indicated proteins. (F) HeLa cells were grown asynchronously or synchronized at G1/S phase by performing a double thymidine block. Cells were then released synchronously back into the cell cycle upon addition of thymidine-free media and cells were collected at various time-points following release. All cell lysates were analyzed by immunoblot for the indicated proteins. (G) HeLa cells were grown asynchronously or synchronized in mitosis by treatment with 100 ng/mL nocodazole for 16 hours. Mitotic-arrested cells were collected using mitotic “shake-off” procedure, washed stringently to removed nocodazole, and released synchronously back into the cell cycle upon addition of nocodazole-free media. Cells were collected at various time-points following release. All cell lysates were analyzed by immunoblot for the indicated proteins.

We found by immunoblot that endogenous AKAP2 abundance is reduced in HCT116 cells arrested in mitosis with nocodazole (**Figure 2C**), but not when cells (HCT116 and RPE1) were arrested in other cell cycle phases **(Figure 2D/E)**. To monitor AKAP2 expression in cycling cells, we released HeLa cells from G1/S phase synchronization following a double-thymidine block. AKAP2 protein levels began to decrease at 10 hours post-release, correlating with the timing of mitosis, as indicated by phospho-H3 S10 staining, and with the peak of PLK1 activation, as measured by phospho-PLK1 T210 and phospho-TCTP S46 (**Figure 2F**). We next monitored AKAP2 expression beginning from mitotic synchronization with nocodazole. AKAP2 protein levels decreased for several hours as cells exited mitosis and re-accumulated 8-10 hours following release, correlating with the start of S-phase, marked by the appearance of Cyclin E1 and Cyclin A (**Figure 2G**). Together, these data demonstrate that AKAP2 is a cell cycle regulated protein whose expression is downregulated in mitosis.

### PLK1 Regulates AKAP2 Abundance in Mitosis

Our proteomic analysis strongly suggested that the downregulation of AKAP2 abundance during mitosis is mediated by PLK1. To test this hypothesis, we arrested cells in mitosis with nocodazole in the presence or absence of small molecule inhibitors targeting PLK1. In HCT116 (**Figure 3A**) and HeLa cells **(Figure S4A)** the decrease in AKAP2 abundance in mitosis was rescued when PLK1 was inhibited with either BI6727 or BI2536. Similarly, the reduction of AKAP2 in mitosis was blocked by treatment with Onvansertib, a third generation PLK1 inhibitor, which is currently under clinical investigation in KRAS mutant colorectal cancer (NCT05593328) **(Figure S4B)**. AKAP2 was also downregulated in a PLK1-dependent manner when cells were synchronized in mitosis using paclitaxel, suggesting that the effect is not related to the microtubule destabilizing effects of nocodazole **(Figure S4C)**. Similarly, AKAP2 protein levels were increased in mitotic cells following transient depletion of PLK1 with siRNA (**Figure 3B**).

**Figure 3.**
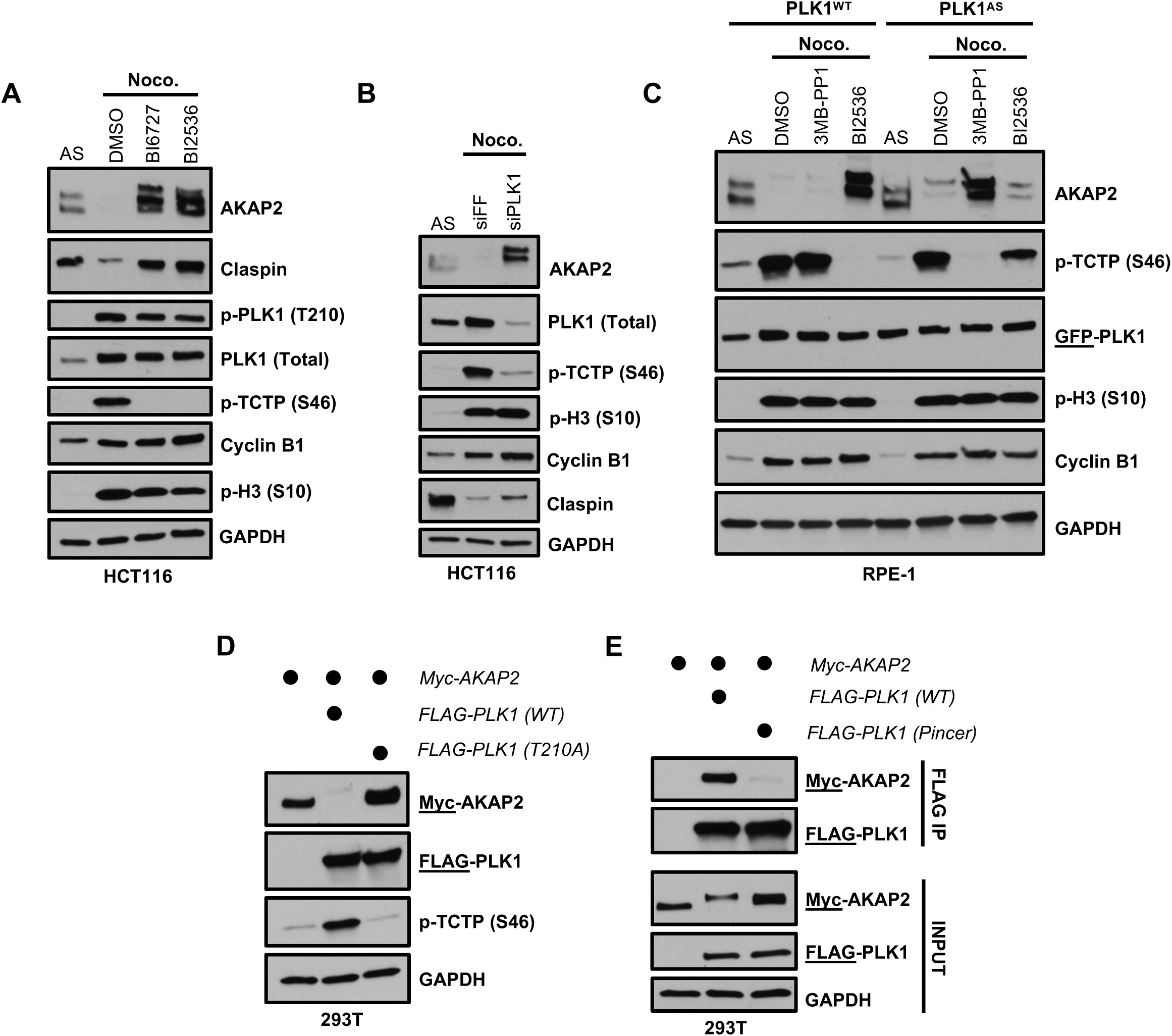
PLK1 Regulates AKAP2 Abundance in Mitosis. (A) HCT116 cells were grown asynchronously or were co-treated with 100 ng/mL nocodazole plus DMSO, 100 nM BI6727, or 100 nM BI2536 for 16 hours. After 16 hours, mitotic-arrested cells were collected using mitotic “shake-off” procedure. All cell lysates were analyzed by immunoblot for the indicated proteins. (B) HCT116 cells were transfected with siRNA targeting PLK1 or firefly luciferase (siFF) as a non-targeting control for 48 hours. During the final 16 hours, cells were treated with 100 ng/mL nocodazole. After 16 hours, mitotic-arrested cells were collected using mitotic “shake-off” procedure. All cell lysates were analyzed by immunoblot for the indicated proteins. (C) Wild-type or PLK1 analog-sensitive (AS) RPE-1 cells were grown asynchronously or were co-treated with 100 ng/mL nocodazole plus DMSO, 10 µM 3MB-PP1, or 100 nM BI2536 for 16 hours. After 16 hours, mitotic-arrested cells were collected using mitotic “shake-off” procedure. All cell lysates were analyzed by immunoblot for the indicated proteins. (D) HEK293T cells were co-transfected with plasmids expressing Myc-AKAP2 together with an empty vector control (lane 1), wild-type FLAG-PLK1 (lane 2), or a catalytically inactive version of FLAG-PLK1 (T210A; lane 3). 24 hours post-transfection, cells were collected and all cell lysates were analyzed by immunoblot for the indicated proteins. (E) HEK293T cells were co-transfected with plasmids expressing Myc-AKAP2 together with an empty vector control (lane 1), wild-type FLAG-PLK1 (lane 2), or a version of FLAG-PLK1 containing a mutation within the polo-box domain (Pincer; lane 3). 24 hours post-transfection, cells were collected and PLK1 was immunoprecipitated using anti-FLAG affinity gel. Eluates were analyzed by immunoblot for the indicated proteins.

To further confirm a direct role for PLK1 in regulating AKAP2 abundance, we utilized a chemical genetic approach using RPE-1 cells in which endogenous PLK1 was replaced with wildtype (WT) or an ATP analog-sensitive (AS) version of PLK1 (Burkard et al. 2007). PLK1-AS has a mutation in the gatekeeper residue which allows for specific PLK1 inhibition using the bulky ATP analog 3MB-PP1. We synchronized PLK1-WT or PLK1-AS cells in mitosis in the presence or absence of either 3MB-PP1 or the small molecule PLK1i BI2536 (**Figure 3C**). As expected, PLK1-WT cells are insensitive to 3MB-PP1, showing no change in PLK1 activity based on the substrate marker p-TCTP (S46) and, consistently, no change in AKAP2 abundance. These cells, however, remain sensitive to the PLK1i BI2536, which caused a decrease in p-TCTP and increase in AKAP2 expression. PLK1-AS cells are sensitive to 3MB-PP1, showing reduced PLK1 activity and a significant increase in AKAP2 abundance. Further, PLK1-AS cells were previously shown to exhibit reduced sensitivity to BI2536 (Burkard et al. 2012) and, consistently, we could not rescue AKAP2 protein levels by BI2536 treatment in PLK1-AS cells. Thus, PLK1 abundance and activity are necessary for the downregulation of AKAP2 in mitosis across multiple cell lines.

We next examined the effects of PLK1 on AKAP2 stability by using ectopic over-expression in HEK293T cells. Levels of exogenously expressed AKAP2 were reduced when co-expressed with wild-type PLK1 (**Figure 3D**). However, no reduction in AKAP2 was seen when co-expressed with a catalytically inactive mutant of PLK1 (PLK1^T210A^). Next, we asked whether PLK1 binds AKAP2 to regulate its abundance. Indeed, wildtype PLK1 interacts with AKAP2, as determined by co-immunoprecipitation when co-expressed exogenously in HEK293T cells (**Figure 3E**). PLK1 uses two key residues in its polo-box domain to bind substrates that have been “primed” by phosphorylation by a separate kinase (Elia et al. 2003). Mutations in these residues (PLK1^Pincer^) abrogate binding to AKAP2, suggesting that the polo-box domain of PLK1 is important for its regulation of AKAP2 (**Figure 3E**). Altogether, these data suggest that PLK1 binds to AKAP2 to regulate its abundance during mitosis.

### SCF**^β^**^TrCP^ Mediates the Proteasomal Degradation of AKAP2 During Mitosis

To understand the mechanism by which AKAP2 expression is reduced during mitosis, we arrested cells in mitosis using nocodazole in the presence of the neddylation inhibitor MLN4924, which inhibits Cullin-RING ligases that depend on neddylation for activity (Soucy et al. 2009), or the proteasome inhibitor bortezomib (**Figure 4A**). The reduction in AKAP2 expression in mitosis was rescued by treatment with each inhibitor, suggesting that AKAP2 is degraded by the ubiquitin-proteasome system, and specifically through the activity of a Cullin-RING ubiquitin ligase.

**Figure 4.**
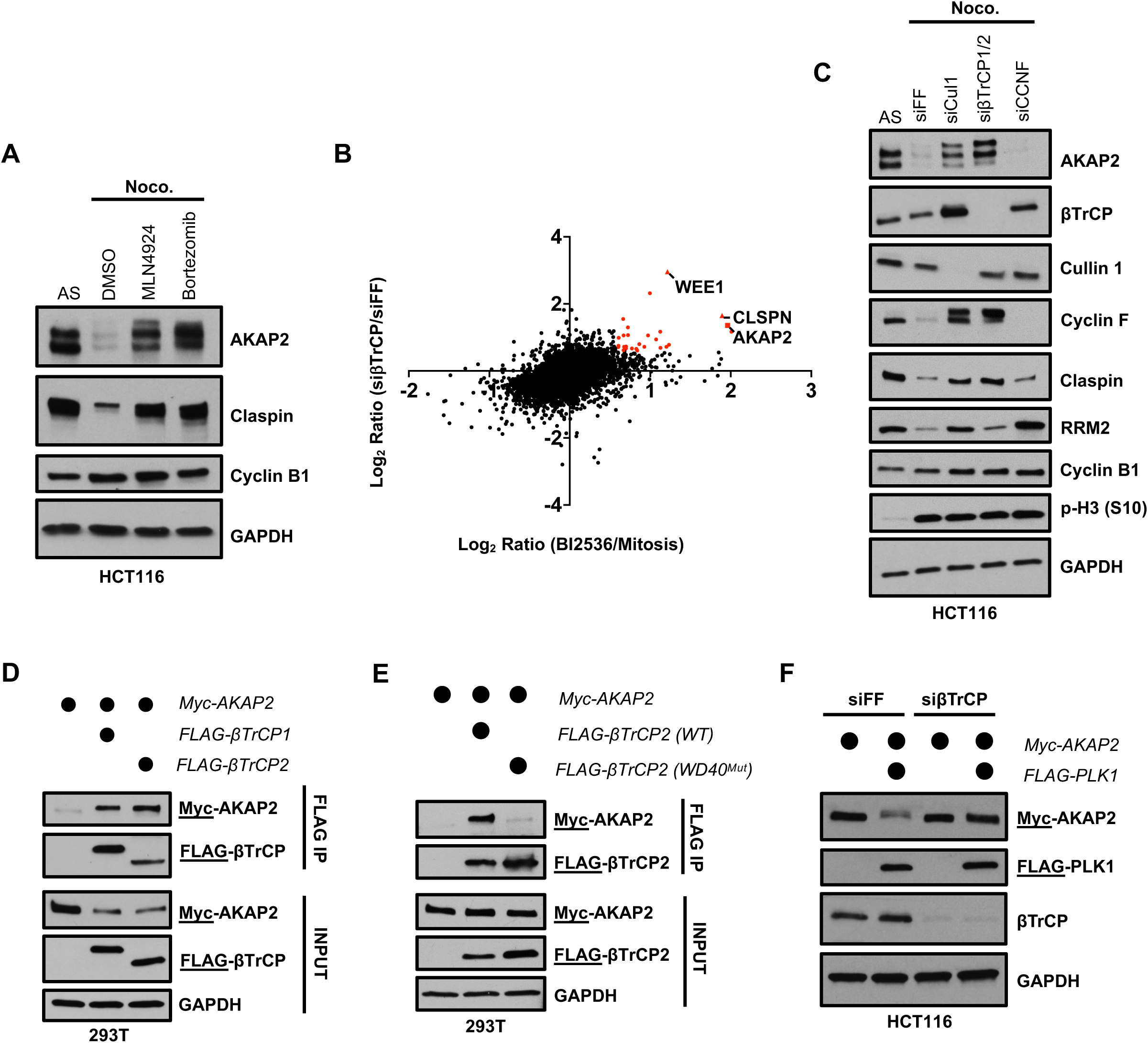
SCF^βTrCP^ Mediates the Proteasomal Degradation of AKAP2 During Mitosis. (A) HCT116 cells were grown asynchronously or were co-treated with 100 ng/mL nocodazole plus DMSO, 100 nM MLN4924, or 100 nM Bortezomib for 16 hours. After 16 hours, mitotic-arrested cells were collected using mitotic “shake-off” procedure. All cell lysates were analyzed by immunoblot for the indicated proteins. (B) Plot showing the Log2 ratio of all proteins identified by mass-spectrometry when comparing samples from BI2536-treated mitotic cells to samples from DMSO-treated mitotic cells (x-axis) and the Log2 ratio of all proteins identified by mass-spectrometry when comparing samples from siβTrCP-treated mitotic cells to samples from siFF-treated mitotic cells (y-axis). Claspin and WEE1 are identified as a positive controls. (C) HCT116 cells were transfected with siRNA targeting Cullin1, βTrCP1/2, CCNF, or firefly luciferase (siFF) as a non-targeting control for 48 hours. During the final 16 hours, cells were treated with 100 ng/mL nocodazole. After 16 hours, mitotic-arrested cells were collected using mitotic “shake-off” procedure. All cell lysates were analyzed by immunoblot for the indicated proteins. (D) HEK293T cells were co-transfected with plasmids expressing Myc-AKAP2 together with an empty vector control (lane 1), wild-type FLAG-βTrCP1 (lane 2), or wild-type FLAG-βTrCP2 (lane 3). 24 hours post-transfection, cells were collected and βTrCP was immunoprecipitated using anti-FLAG affinity gel. Eluates were analyzed by immunoblot for the indicated proteins. (E) HEK293T cells were co-transfected with plasmids expressing Myc-AKAP2 together with an empty vector control (lane 1), wild-type FLAG-βTrCP2 (lane 2), or a version of FLAG-βTrCP2 with a mutation in the substrate binding domain (WD40^Mut^; lane 3). 24 hours post-transfection, cells were collected and βTrCP was immunoprecipitated using anti-FLAG affinity gel. Eluates were analyzed by immunoblot for the indicated proteins. (F) HCT116 cells were transfected with siRNA targeting βTrCP1/2 or firefly luciferase (siFF) as a non-targeting control for 24 hours. After 24, hours, cells were co-transfected with plasmids expressing Myc-AKAP2 together with an empty vector control (lanes 1,3), or with wild-type FLAG-PLK1 (lanes 2,4) and cultured for an additional 24 hours. Then, cells were collected and all cell lysates were analyzed by immunoblot for the indicated proteins.

To determine the role of βTrCP and Cyclin F in shaping the mitotic proteome, and their potential role in AKAP2 degradation, we transiently depleted βTrCP or Cyclin F using siRNA, arrested cells in mitosis, and performed label-free, quantitative proteomics **(Figure S5A-C and Table S3)**.

Several of the top hits in the βTrCP depleted samples were established substrates, including WEE1, USP37, and Claspin **(Figure S5B)**. AKAP2 was also among these top hits, showing a 2.6-fold increase in expression in mitotic cells depleted of βTrCP compared to our non-targeting control **(Figure S5B)**. When comparing the data from our PLK1i proteomics with our βTrCP siRNA proteomics, AKAP2 scored among the most up-regulated proteins in both datasets, strongly suggesting that its degradation is mediated along the PLK1-βTrCP signaling axis, similar to WEE1 and Claspin (**Figure 4B**). Indeed, siRNA knockdown followed by immunoblot confirmed that AKAP2 degradation is dependent on Cullin 1, the scaffold protein utilized by all F-box proteins, and βTrCP, but not Cyclin F (**Figure 4C**). Consistently, we were able to detect an interaction between AKAP2 and both βTrCP1 and βTrCP2 by co-IP following ectopic expression in HEK293T cells (**Figure 4D**). Moreover, mutations in the WD40 repeats of βTrCP1 (βTrCP1^WD40Mut^) and βTrCP2 (βTrCP2^WD40Mut^) that disrupt binding to substrates (Wu et al. 2003; Kruiswijk et al. 2012) prevent their interaction with AKAP2 (**Figure 4E** **and Figure S5D)**.

Next, we determined whether βTrCP cooperates with PLK1 to promote AKAP2 degradation. We expressed AKAP2 together with an empty vector control or with PLK1 in HCT116 cells treated with siRNA targeting firefly luciferase (non-targeting control) or βTrCP (**Figure 4F**). Whereas AKAP2 abundance decreased following PLK1 expression in control cells, this effect was abrogated in cells depleted of βTrCP. These data further support the notion that SCF^βTrCP^ is the E3 ubiquitin ligase responsible for the proteasomal degradation of AKAP2 during mitosis.

### SCF**^β^**^TrCP^ Utilizes a Non-Canonical Degron to Bind and Degrade AKAP2

The canonical degron sequence found in βTrCP substrates is DSGxx(x)S, in which both serine residues are typically required to be phosphorylated for efficient βTrCP binding. However, many substrates with variable βTrCP degrons have been reported, such as CDC25A and CDC25B (Kanemori et al. 2005). AKAP2 does not have a fully canonical βTrCP degron, however it does contain 3 ‘DSG’ minimal sequence motifs (**Figure 5A**). We examined binding to βTrCP of 3 separate AKAP2 mutants in which the DSG motifs were individually disrupted by mutating each residue within the motif to alanine (referred to as DSG1, DSG2, or DSG3). Notably, DSG1 is not conserved across species whereas DSG2 and DSG3 are well-conserved (**Figure 5B** **and Figure S6A/B)**. AKAP2^WT^, AKAP2^DSG1^ and AKAP2^DSG3^ all similarly co-immunoprecipitated with βTrCP (**Figure 5C**). However, AKAP2^DSG2^ (amino acids 472-474) was unable to interact with βTrCP, suggesting that DSG2 is likely the functional degron.

**Figure 5.**
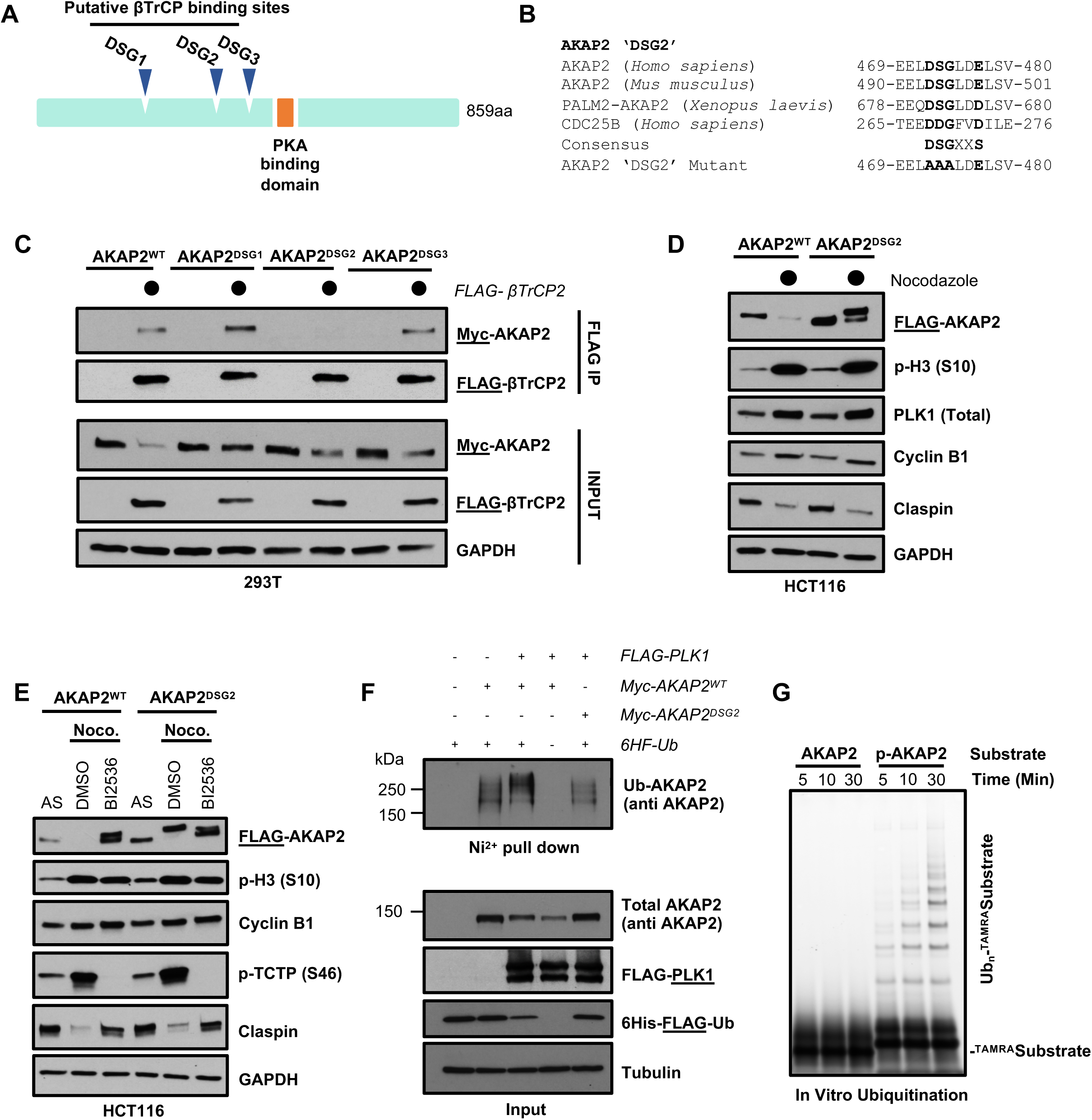
βTrCP Utilizes a Non-Canonical Degron to Degrade AKAP2 During Mitosis. (A) Schematic depicting AKAP2 showing the PKA binding domain and relative positions of the three putative βTrCP binding sites in AKAP2. The corresponding amino acids for each motif are 382-387 (‘DSG1’), 472-477 (’DSG2’), and 502-507 (‘DSG3’). (B) Sequence alignment of the ‘DSG2’ motif in *Homo sapiens, Mus musculus,* and *Xenopus laevis.* The CDC25B degron sequence for βTrCP is also shown, demonstrating a known example which is divergent from the canonical ‘DSGxxS’ sequence. The sequence of the ‘DSG2’ motif AKAP2 mutant used in future experiments is shown below. (C) HEK293T cells were co-transfected with plasmids expressing Myc-AKAP2 (WT or ‘DSG’ degron mutants) together with an empty vector control or FLAG-βTrCP2. 24 hours post-transfection, cells were collected and βTrCP was immunoprecipitated using anti-FLAG affinity gel. Eluates were analyzed by immunoblot for the indicated proteins. (D) HCT116 cells stably expressing FLAG-AKAP2^WT^ or FLAG-AKAP2^DSG2^ were grown asynchronously or were treated with 100 ng/mL nocodazole for 16 hours to synchronize cells in mitosis. After 16 hours, mitotic-arrested cells were collected using mitotic “shake-off” procedure. All cell lysates were analyzed by immunoblot for the indicated proteins. (E) HCT116 cells stably expressing FLAG-AKAP2^WT^ or FLAG-AKAP2^DSG2^ were grown asynchronously or were co-treated with 100 ng/mL nocodazole plus DMSO or 100 nM BI2536 for 16 hours. After 16 hours, mitotic-arrested cells were collected using mitotic “shake-off” procedure. All cell lysates were analyzed by immunoblot for the indicated proteins. (F) HEK293T cells were co-transfected with the indicated plasmids for 48 hours. After 48 hours, cells were collected and lysed under denaturing conditions. Ubiquitinated proteins were captured using Ni-NTA pulldown. The ubiquitination status of AKAP2 was analyzed by immunoblot. Cells were treated with 20 μM MG132 and 20 μM PR619 for 4 hours prior to collecting cells. (G) In vitro ubiquitination reactions using TAMRA-labeled AKAP2 WT or phospho-peptide as a substrate, monitored by fluorescent scanning of an SDS-PAGE gel. Representative of n=4 independent experiments.

Next, we tested whether the DSG2 mutant is stabilized in mitosis. To do so, we generated HCT116 cells stably expressing either FLAG-AKAP2^WT^ or FLAG-AKAP2^DSG2^. In asynchronous cells, FLAG-AKAP2^DSG2^ was more abundant than FLAG-AKAP2^WT^, consistent with the lack of binding to βTrCP **(Figure S6C)**. Significantly, similar to endogenous AKAP2, FLAG-AKAP2^WT^ was degraded in mitosis, whereas FLAG-AKAP2^DSG2^ was resistant to degradation (**Figure 5D**). Other βTrCP substrates, such as Claspin, were degraded normally in cells expressing FLAG-AKAP2^DSG2^. Stabilized AKAP2 does not interfere with the cell’s ability to arrest in mitosis, as indicated by equal levels of Cyclin B1 and phospho-H3 S10. We also engineered HCT116 cells in which expression of AKAP2^WT^ or AKAP2^DSG2^ was under the control of a doxycycline-inducible promoter. To monitor AKAP2 stability, we arrested these cells in mitosis in the presence of doxycycline to induce AKAP2 expression **(Figure S6D)**. Using the same concentration of doxycycline, AKAP2^DSG2^ expression was higher than that of AKAP2^WT^ suggesting that AKAP2^DSG2^ is resistant to degradation in mitosis.

The role of PLK1 in promoting the degradation of βTrCP substrates is to phosphorylate the target at one of the serine residues within the βTrCP degron sequence to promote the interaction between the E3 ligase and its substrate. Thus, AKAP2^DSG2^ should be unaffected by PLK1 inhibition. To test this hypothesis, we arrested our FLAG-AKAP2^WT^ or FLAG-AKAP2^DSG2^ expressing HCT116 cells in mitosis with or without PLK1 inhibition using BI2536 (**Figure 5E**). FLAG-AKAP2^WT^ was degraded in mitosis and this effect was reversed upon inhibition of PLK1. Conversely, FLAG-AKAP2^DSG2^ was not degraded in mitosis and showed no change in abundance when PLK1 was inhibited with BI2536. Interestingly, although abundance of AKAP2^DSG2^ was unaffected by PLK1 inhibition, there was a slight electrophoretic shift of the protein, suggesting that PLK1 may phosphorylate AKAP2 at additional, unidentified sites. Consistently, AKAP2^DSG2^ is still able to interact with PLK1 by co-IP **(Figure S6E)**. When co-expressed ectopically with PLK1 in HEK293T cells, AKAP2^WT^ is degraded, whereas AKAP2^DSG2^ is not **(Figure S6E; see input lanes)**. However, PLK1 expression results in a slower migrating form of AKAP2^DSG2^, further supporting the notion that PLK1 phosphorylates AKAP2 at additional sites that are independent of its role in promoting degradation.

Next, we assessed the ability of PLK1 to promote the ubiquitination of AKAP2^WT^ or AKAP2^DSG2^ using an *in vivo* ubiquitination assay following ectopic expression in HEK293T cells (**Figure 5F**). These cells were transfected with Myc-tagged AKAP2 (WT or DSG2) and 6xHIS-FLAG-ubiquitin in combination with FLAG-PLK1. To increase total protein abundance and to prevent de-ubiquitination, cells were treated with the proteasome inhibitor MG132 and the pan-DUB inhibitor PR-619 for 4 hours prior to harvesting. Cells were then lysed under denaturing conditions and subjected to ubiquitin purification by pull-down on Ni^2+^-NTA resin. Expression of PLK1 increased the polyubiquitination of AKAP2^WT^, as indicated by the slower migrating species of AKAP2 observed by SDS-PAGE (compare lane 2 vs. lane 3). However, even in the presence of PLK1, AKAP2^DSG2^ displayed less ubiquitination than AKAP2^WT^ expressed alone (compare lane 2 vs. lane 5). This suggests that an intact DSG2 motif is important for the ubiquitination of AKAP2.

Finally, we reconstituted the ubiquitination of AKAP2 *in vitro* (**Figure 5G**). Purified components of the SCF^βTrCP^ complex were mixed with ubiquitin, E1 and E2 enzymes, and a fluorescently labeled AKAP2 peptide containing the DSG2 motif. Notably, ubiquitination of the peptide was only observed when the serine residue within the DSG2 degron was phosphorylated. Unphosphorylated and phosphorylated β-catenin peptides, which are similarly regulated by SCF^βTrCP^ served as a positive control **(Figure S6F)**. Together, these data suggest that PLK1 phosphorylates AKAP2 within its second DSG-degron motif, which allows βTrCP to bind to and ubiquitinate AKAP2.

### The AKAP2 Interactome Reveals a Role for AKAP2 in Regulating Proliferative Signaling Networks

AKAP2 is poorly studied and has no reported roles in cell cycle regulation. We sought to understand the biological significance of AKAP2 in cell cycle and proliferation. To do so, we defined the AKAP2 interactome by performing immunoprecipitation mass-spectrometry (IP-MS) analysis of triplicate samples using exogenously expressed Myc-AKAP2 in HEK293T cells (**Figure 6A** **and Table S4)**. We recovered the entire SCF^βTrCP^ complex as well as both the catalytic and regulatory subunits of PKA, which are known interactors of the AKAP proteins (**Figure 6B** **and Figure S7A)**. In addition, many of the top AKAP2 interactors are involved in coordinating various signaling transduction pathways, including multiple components of the protein phosphatase 2A complex as well as the striatin-interacting phosphatase and kinase complex (STRIPAK). Consistently, GO analysis on the AKAP2 interactors shows enrichment for processes including “Cell Cycle,” “G2/M Transition,” “Intracellular signaling by second messengers,” and “Cell division” (**Figure 6C** **and Table S2).** We validated several AKAP2 interactors using immunoblot including multiple components of the STRIPAK complex (PDCD10, STK24, STK26), as well as the adaptor protein GRB2 (**Figure 6D**). PKA signaling and the STRIPAK complex play established role in Hippo signaling and GRB2 is an intermediary between receptor tyrosine kinases and activation of the Ras-MAPK pathway (Kong et al. 2019; Rozakis-Adcock et al. 1993; Yu et al. 2013).

**Figure 6.**
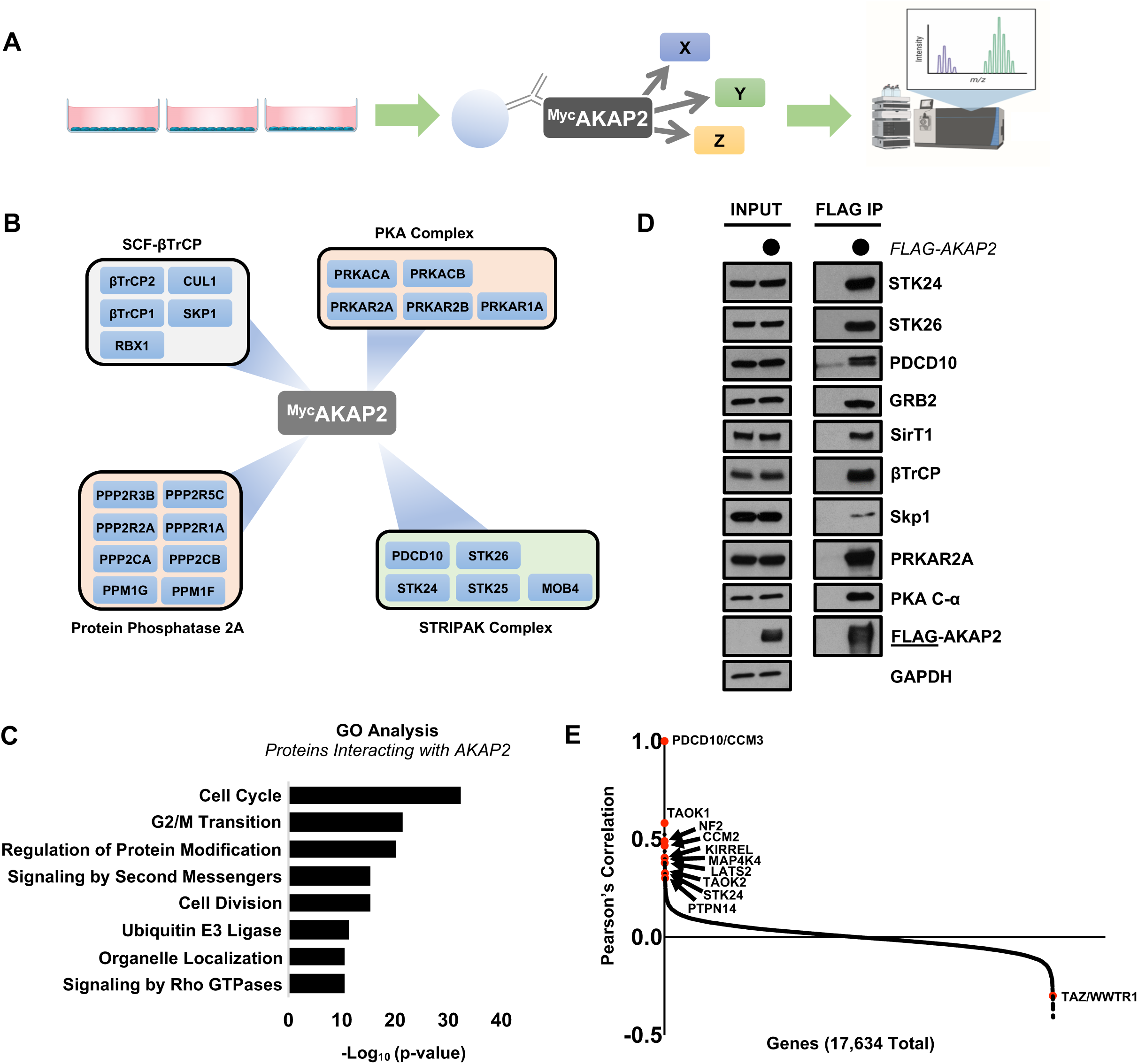
The AKAP2 Interactome Reveals a Role for AKAP2 in Regulating Proliferative Signaling Networks. (A) AKAP2 IP-MS Schematic. Myc-AKAP2 was transfected into HEK293T cells in triplicate. AKAP2 was immunoprecipitated using anti-Myc affinity gel and samples were analyzed by LC-MS/MS. (B) AKAP2 interactome showing strongest interactors and the complexes they function in as identified by IP-MS. (C) GO analysis highlighting top biological processes enriched among the strongest AKAP2 interacting proteins. (D) HEK293T cells were transfected with Myc-AKAP2 or an EV control for 24 hours. After 24 hours, AKAP2 was immunoprecipitated using anti-Myc affinity gel and eluates were analyzed by immunoblot for the indicated proteins. (E) DepMap analysis of PDCD10 showing that it is highly correlated with genes that function within the Hippo Signaling Pathway.

To obtain further insight into the physiological processes the AKAP2 interactors regulate, we relied on analysis of the Project Achilles Cancer Dependency Map (DepMap). The DepMap uses CRISPR/Cas9 loss-of-function screens to provide a fitness score for each gene based on their impact on cell proliferation/viability. Importantly, genes whose fitness scores are highly correlated with each other are often physically or genetically linked to one another and may function within the same pathways (Hart et al. 2015; Enrico et al. 2021). PDCD10, the strongest AKAP2 interactor, is highly correlated with multiple genes functioning within the Hippo tumor suppressor network (**Figure 6E**). GRB2 is highly correlated with genes that coordinate the MAPK and AKT signaling pathways **(Figure S7B)**. Together, this suggests that AKAP2 regulates various proliferative networks and that PLK1-mediated degradation of AKAP2 might couple cell cycle progression and cell division with proliferative decisions and cell growth.

## Discussion

Ubiquitin-mediated proteolysis is critical for proliferation and, in combination with transcriptional dynamics, facilitates the widespread proteome remodeling which accompanies progression through the cell cycle (Potapova et al. 2006). This is most well-established at the mitosis-to-G1 transition, when the APC/C ubiquitin ligase facilitates large-scale changes in the protein landscape by driving the degradation of many dozen substrates (Davey and Morgan 2016; Franks et al. 2020). The SCF-family of E3 ligases play similarly important and evolutionarily conserved roles in the cell cycle. Skp1, an adaptor protein for the SCF, is required for both the G1/S and G2/M transitions (Bai et al. 1996; Connelly and Hieter 1996). Additionally, yeast mutants defective for the SCF components Cdc4, Cdc34 and Cdc53 all arrest in G1-phase (Schwob et al. 1994) because of their inability to degrade the CDK inhibitor Sic1 (Connelly and Hieter 1996; Bai et al. 1996). Likewise, SCF-mediated degradation of the CDK inhibitor p27 in higher eukaryotes is essential for entry into the S-phase of the cell cycle (Carrano et al. 1999; Kossatz et al. 2004). Despite the widespread role of ubiquitin signaling in sculpting cell cycle proteomes, the overall contribution of protein ubiquitination and degradation at G2/M has remained less well-characterized.

Significantly, our proteomic analysis of mitotically arrested cells found that over 200 proteins are downregulated in mitosis, several of which are previously established substrates of SCF ubiquitin ligases. Thus, the magnitude of protein degradation that occurs at G2/M is similar to what is seen at mitotic exit. Many of these proteins are also likely to oscillate transcriptionally, since many cell cycle ubiquitin substrates are also controlled by cell cycle transcriptional dynamics (Franks et al. 2020). In addition, our approach is likely to have captured proteins degraded at different points prior to mitotic entry. It will be interesting in the future to assess specific degradative timing patterns for these diverse proteins.

Given previous reports showing that PLK1 drives the degradation of some SCF^βTrCP^ substrates at G2/M (Hansen et al. 2004; Burrows et al. 2012; Watanabe et al. 2004; Peschiaroli et al. 2006; Mailand et al. 2006; Mamely et al. 2006; Seki et al. 2008; Moshe et al. 2004; Chan et al. 2008), we further tested the contribution of PLK1 in shaping the mitotic proteome. Remarkably, we found that the abundance of over 200 proteins is increased in mitosis following PLK1 inhibition. Moreover, more than half of the proteins (∼56%) that we found to be downregulated in mitosis were subsequently rescued by PLK1 inhibition. Importantly, this suggests that PLK1 mediates a specific and widespread proteolytic program which we predict is confined to the timing of PLK1 activation in the G2/M phase. Interestingly, while many PLK1-regulated proteins are involved in cell cycle regulation, gene ontology analysis revealed that several proteins involved in transcriptional and ribosomal RNA processes are also sensitive to PLK1 inhibition. We speculate that PLK1-mediated ubiquitin signaling contributes to the transcriptional and translational repression associated with mitosis (Palozola et al. 2017; Tanenbaum et al. 2015).

Prior to this study, PLK1 has exclusively been shown to regulate the degradation of SCF^βTrCP^ substrates at G2/M. Here, we found that the degradation of SCF^Cyclin^ ^F^ substrates during mitosis is also PLK1-dependent, suggesting that PLK1 coordinates protein degradation through at least two E3 ubiquitin ligases. To connect newly identified PLK1-dependent substrates with their respective E3 ligases, we performed proteomics analysis of mitotic cells depleted of either βTrCP or Cyclin F. However, the proteins upregulated by knockdown of these two ligases do not account for all of the proteins sensitive to PLK1 inhibition. Thus, there are likely yet-to-be-identified E3 ubiquitin ligases that cooperate with PLK1 to promote protein degradation at G2/M. There are also likely PLK1-independent protein degradation events that remain to be explored.

Generally, substrates of SCF ubiquitin ligases require phosphorylation within defined sequences, termed phospho-degrons, to promote binding to the ligase and subsequent degradation (Skowyra et al. 1997; Ang and Wade Harper 2005). Although the need for phosphorylation of some substrates has been suggested (D’Angiolella et al. 2012), Cyclin F is unique in that it does not directly recognize phospho-degrons. This is illustrated by the fact that Cyclin F substrates can be ubiquitinated *in vitro* without the need for phosphorylation (Enrico et al. 2021). Although we have not elucidated the full mechanism, our data suggest that PLK1 regulates Cyclin F-dependent degradation at the ligase-level by promoting the stability of the Cyclin F protein, rather than through control of each individual substrate. There are no reported interactions between PLK1 and Cyclin F substrates that function to promote their proteolysis. We speculate that this mode of regulation could allow PLK1 activity, which changes dynamically during cell cycle and in response to DNA damage, to cooperate with other signaling inputs to control distinct sets of substrates at particular times or in response to specific stresses. Moreover, since some Cyclin F substrates are degraded earlier in the cell cycle, we predict that other factors, such as localization, could define the ability of Cyclin F to bind and ubiquitinate substrates. Interestingly, we have previously shown that Cyclin F is stabilized by AKT earlier in the cell cycle (Choudhury et al. 2017). There may therefore be cooperation between AKT and PLK1 to regulate Cyclin F stability throughout the cell cycle. Collectively this suggests that the regulation of Cyclin F substrates depends on ligase-level control. This is in contrast with most other SCF-ligases, and with βTrCP in particular, where the destruction of substrates is dependent on substrate-level control. Altogether, this could represent a way to regulate the ordering of substrate destruction, which is likely to be important for proper cell cycle progression.

We identified the A-kinase anchor protein 2 (AKAP2) as a novel cell cycle regulated protein whose mitotic degradation is regulated along the PLK1/βTrCP signaling axis. The role of AKAP family proteins is to scaffold protein kinase A (PKA) in distinct areas of the cell. This helps to create localized PKA signaling hubs and allows for specificity, acceleration, and amplification of PKA-dependent responses (Greenwald and Saucerman 2011). Therefore, AKAP proteins generally bind to a subcellular structure, the regulatory subunit of PKA, and a multitude of proteins involved in signal transduction. AKAP2 has been reported to bind actin (Dong et al. 1998); however, only a few other interactors of AKAP2 have been reported.

PKA plays a well-established role in cell division, although this has been most well-established in meiosis (Grieco et al. 1996; Maller and Krebs 1977). Several AKAPs are reported to contribute to processes of cell cycle regulation including regulation of the G1/S transition, the completion of cytokinesis, and chromatin condensation (Han et al. 2015). However, AKAP2 has no reported role in cell cycle regulation. In addition, to the best of our knowledge, no other AKAP has been shown to oscillate in abundance during cell cycle progression. The cell cycle regulation of AKAP2 protein expression and the AKAP2 interactome suggest that AKAP2 contributes to cell cycle progression. The highest confidence AKAP2 interactors function in proliferative signaling networks including the MAPK, AKT, and Hippo signaling pathways. This suggests that AKAP2, and its mitotic degradation, could integrate proliferative signaling networks during the cell cycle. Since AKAP2 binds and localizes to actin (Dong et al. 1998), we propose that it tethers PKA to the cytoskeleton during interphase to facilitate phosphorylation of yet-to-be-identified substrates. During G2/M, AKAP2 is phosphorylated by PLK1 and degraded by SCF^βTrCP^. As cells enter mitosis, when AKAP2 protein is no longer present, we hypothesize that this allows for the separation of PKA from its AKAP2-dependent substrates, leading to substrate de-phosphorylation (**Figure 7**). Determining the direct PKA-AKAP2 substrates and the downstream physiological consequences of their regulation will be important for understanding the role of AKAP2 in the cell cycle.

**Figure 7.**
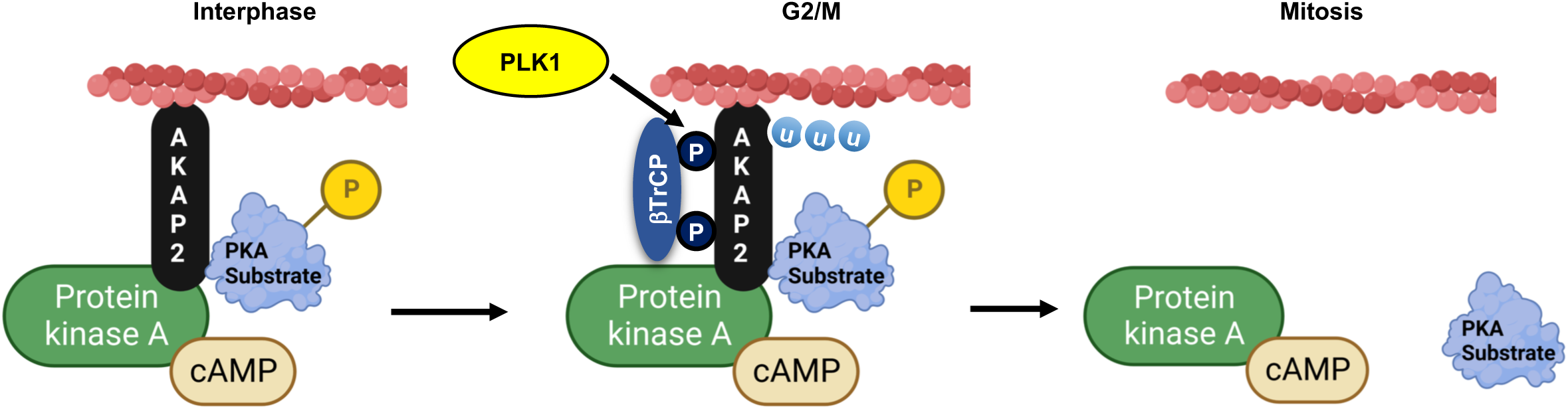
Proposed Model for PLK1/βTrCP-Mediated Degradation of AKAP2. (A) During interphase, AKAP2 is expressed and is predicted to scaffold PKA with specific substrates leading to their phosphorylation. Based on our IP-MS data as well as previous reports, AKAP2 is depicted as being localized to the actin cytoskeleton. However, it is possible that AKAP2 is also present at other subcellular locations. As cells approach G2/M, PLK1 phosphorylates AKAP2 which allows **β**TrCP to bind to and ubiquitinate AKAP2. Then, by the time cells reach mitosis AKAP2 is degraded. It is predicted that this then causes the dissociation between PKA and it’s AKAP2-dependent substrates, which could allow for their dephosphorylation. The specific substrates that AKAP2 directs PKA to are still undetermined.

Our data could have clinical implications since PLK1 is often over-expressed in various cancers (Liu et al. 2017) and is the target of small molecule inhibitors under investigation for clinical use (Iliaki et al. 2021). The third generation PLK1 inhibitor onvansertib generated promising clinical trial data in patients with metastatic colorectal cancer harboring activating KRAS mutations (Lenz et al. 2022). Typically, sensitivity and resistance to targeted therapies is studied by examining alterations in the phospho-proteome, transcriptome, or kinome. Given our data, it is interesting to consider the possibility that remodeling of the proteome through changes in protein degradation will also contribute to both therapeutic responses and treatment resistance. The clinical use of PLK1 inhibitors illustrates the importance of further investigating this possibility.

Many of the proteins which we identified as being controlled by PLK1 are linked to tumorigenesis. Therefore, failure to degrade these proteins in response to PLK1 targeted therapies could provide an unanticipated avenue by which cells adapt to therapy-induced proliferative arrest. While we validated several PLK1-dependent substrates in multiple cell lines, a global analysis across multiple cell lines will be beneficial in the future for understanding how the proteome is remodeled by PLK1 and PLK1 inhibitors in different cell and cancer types. This is especially true if, as we predict, different signaling inputs cooperate with PLK1 to contribute to the destruction of specific sets of substrates. This understanding could nominate therapeutic situations in which PLK1 inhibitors might be most beneficial.

## Experimental Procedures

### Cell Lines, Cell Culture, and Drug Treatments

Cell lines used in this study include: HCT116 (ATCC), HeLa (ATCC), RPE-1-hTRET (generously provided by Peter Jackson, Stanford University), HEK293T (ATCC), T47D (ATCC), MDA-MB-231 (ATCC), HeLa sgCntrl and HeLa sgCCNF (Choudhury et al. 2016), and RPE-1 EGFP-PLK1 WT and RPE-1 EGFP-PLK1 AS (generously provided by Prasad Jallepalli, MSKCC) (Burkard et al. 2007). These cell lines were cultured in high glucose containing Dulbecco’s Modified Eagle’s Medium (DMEM; Gibco; cat. #11995) supplemented with 10% fetal bovine serum (FBS; VWR) and 1% penicillin/streptomycin (Gibco). Additional cell lines generated and used in this study include: HCT116 pGenLenti AKAP2 WT, HCT116 pGenLenti AKAP2 DSG2, HCT116 pIND20 EV, HCT116 pIND20 AKAP2 WT, and HCT116 pIND20 AKAP2 DSG2. These cell lines were cultured in high glucose containing DMEM supplemented with 10% FBS without antibiotics. All cells were incubated at 37°C, 5.0% CO_2_.

DNA transfection experiments were performed in HEK293T or HCT116 cells. Transfections were performed using Polyjet (SignaGen; cat. #SL100688) transfection reagent according to the manufacturer’s protocol. Transfected cells were cultured for 24 hours prior to analysis. siRNA transfections were performed using RNAiMAX (Thermo Fisher Scientific; cat. #13778075) transfection reagent according to the manufacturer’s protocol. All siRNA transfections were performed at a final siRNA concentration of 20 nM. siRNA information is found in **Table S5.**

For the G1/S-block and release experiment performed in Figure 2, cells were synchronized using a double thymidine block and release protocol. Briefly, cells were treated with 2 mM thymidine for 16 hours. The cells were then extensively washed to remove thymidine and were cultured in thymidine-free media for 8 hours. After 8 hours, cells were treated with 2 mM thymidine for an additional 16 hours. After the 2^nd^ thymidine block, cells were extensively washed and cultured in thymidine-free media for indicated amounts of time. The mitotic block and release experiment in Figure 2 was performed using a nocodazole block and release protocol. Briefly, cells were treated with 100 ng/mL nocodazole for 16 hours. After 16 hours, mitotic-arrested cells were isolated using mitotic “shake-off” procedure. These cells were extensively washed to remove nocodazole and cells were re-plated and cultured in nocodazole-free media for indicated amounts of time.

Information on all other drugs/compounds used in this study can be found in **Table S5**.

### Cell Lysis and immunoblotting

Cells were lysed on ice for 15 minutes in NETN lysis buffer (20 mM Tris pH 8.0, 100 mM NaCl, 0.5 mM EDTA, 0.5% NP40) supplemented with 10 µg/mL aprotinin, 10 µg/mL leupeptin, 10 µg/mL pepstatin A, 1 mM sodium orthovanadate, 1 mM sodium fluoride, and 1 mM AEBSF (4-[two aminoethyl] benzenesulfonyl fluoride). Following incubation, cell lysates were centrifuged at 20,000×*g* in a benchtop microcentrifuge at 4°C for 10 minutes. Protein concentration was determined by Bradford assay (Bio-Rad; cat. #5000006) and samples were normalized and prepared by boiling in Laemmli buffer (LB) at 95°C for 5 minutes on a dry water-bath. Equal amounts of protein were separated by electrophoresis on 4-15% TGX (Bio-Rad) stain-free gels at 150-230V and then transferred to nitrocellulose membranes at 0.2A for 80 minutes using the wet transfer method. After transfer, membranes were blocked in 5% non-fat dry milk (Biorad; cat. #1706404) diluted in 1x TBS-T (137 mM NaCl, 2.7 mM KCl, 25 mM Tris pH 7.6, 1% Tween-20). The majority of the primary antibody incubations were carried out overnight, while rocking, at 4°C. Primary antibody incubations for PLK1 and Cyclin B1 blots were carried out for 30 minutes, while rocking, at room temperature. All HRP-conjugated antibody incubations were carried out for 1 hr, while rocking, at room temperature. 1x TBS-T was used for all wash steps. Protein abundance was visualized by chemiluminescence using Pierce ECL (Thermo Fisher Scientific; cat. #32106). A detailed list of information for primary and secondary antibodies is provided in **Table S5.**

### Sample Preparation for PLK1i and siRNA Mass Spectrometry

For the PLK1 mass spectrometry screen, HCT116 cells were seeded in triplicate into 60 mm plates 24 Hr prior to cell treatments. The following day, cells were left un-treated (asynchronous), or co-treated with 100 ng/mL nocodazole plus DMSO (0.01%), BI2536 (100 nM), or BI6727 (100 nM). After 16 hours, cells were harvested using mitotic “shake-off” procedure. Isolated cells were pelleted by centrifugation at 200×*g* for 8 minutes at 4°C using a tabletop centrifuge. Prior to cell lysis, the cell pellets were washed 4x with 1x dPBS. Cells were lysed on ice for 20 minutes using urea lysis buffer (8M Urea, 50 mM Tris-HCl pH 8.0, 75 mM NaCl, 1 mM EDTA) supplemented with 10 µg/mL aprotinin, 10 µg/mL leupeptin, 10 µg/mL pepstatin A, 1 mM sodium orthovanadate, 1 mM sodium fluoride, and 1 mM AEBSF (4-[two aminoethyl] benzenesulfonyl fluoride). Samples were snap frozen 2x using liquid nitrogen and cell lysates were then clarified by centrifugation at maximum speed (20,000×*g*) for 15 minutes at 4°C using a benchtop microcentrifuge. Protein concentration was determined using Bradford Assay and lysates were analyzed by mass spectrometry, as described.

For the siRNA mass spectrometry, HCT116 cells were seeded in triplicate into 60 mm plates 24 Hr prior to cell treatments. The following day, cells were transfected with siRNAs targeting firefly luciferase (negative control), βTrCP1/2, or Cyclin F at a total concentration of 20 nM siRNA. Cells were cultured with siRNA for a total of 48 hours and were treated with 100 ng/mL nocodazole for the final 16 hours to arrest cells in mitosis. Cells were harvested using mitotic “shake-off” and lysed as described above for the PLK1i mass-spectrometry screen. Protein concentration was determined using BCA Assay (ThermoFisher; cat. PI-23227) and lysates were analyzed by mass spectrometry, as described below.

### Mass Spectrometry

#### MS sample preparation for global, quantitative proteomics

For each sample, 25ug of protein lysate was reduced with 5mM dithithreitol (DTT; Pierce) at 37°C for 45 min and alkylated with 15mM iodoacetamide (IAA; Pierce) for 45 min at room temperature. Samples were then precipitated by adding six volumes of ice-cold acetone and incubating at -20°C overnight. The following day, samples were centrifuged at 15000xg for 15 min at 4°C, supernatant was removed, and the pellet was washed with 100ul cold acetone. Pellets were air dried at room temperature for 10 min before being resuspended in 100ul 50mM ammonium bicarbonate, pH 8. Samples were then subjected to digestion with LysC (Wako) at 37°C for 2h and trypsin (Promega) overnight at 37°C at a 1:50 enzyme:protein ratio. The resulting peptides were acidified to 0.5% trifluoroacetic acid (TFA; Pierce) and desalted using Thermo desalting spin columns. Eluates were dried via vacuum centrifugation and peptide concentration was determined via Pierce Quantitative Fluorometric Assay and all samples were normalized to 0.25 ug/ul and subjected to LC-MS/MS analysis.

#### LC-MS/MS for global, quantitative proteomics

Samples were analyzed in a randomized order by LC-MS/MS using an Ultimate 3000 coupled to an Exploris 480 mass spectrometer (Thermo Scientific). The pooled sample was analyzed before and after the sample set. Samples were injected onto an IonOpticks Aurora series 2 C18 column (75 μm id × 15 cm, 1.6 μm particle size; IonOpticks) and separated over a 200 min method. The gradient for separation consisted of 3-41% mobile phase B at a 250 nl/min flow rate, where mobile phase A was 0.1% formic acid (Pierce) in LC-MS grade water (Fisher) and mobile phase B consisted of 0.1% formic acid in 80% acetonitrile (ACN). Exploris 480 was operated in product ion scan mode for Data Independent Acquisition (DIA).

A full MS scan (m/z 350-1650) was collected; resolution was set to 120,000 with a maximum injection time of 20 ms and automatic gain control (AGC) target of 300%. Following the full MS scan, a product ion scan was collected (30,000 resolution) and consisted of stepped higher collision dissociation (HCD) set to 25.5, 27, 30; AGC target set to 3000%; maximum injection time set to 55 ms; variable precursor isolation windows from 350-1650 m/z.

#### Data analysis for global quantitative proteomics

Raw data files were processed using Spectronaut (v15.7.220308.50606; Biognosys) and searched against the Uniprot reviewed human database (UP000005640, containing 20,396 entries, downloaded March 2021) and the MaxQuant common contaminants database (246 entries). The following settings were used: enzyme specificity set to trypsin, up to two missed cleavages allowed, cysteine carbamidomethylation set as a fixed modification, methionine oxidation and N-terminal acetylation set as variable modifications. Precision iRT calibration was enabled. A false discovery rate (FDR) of 1% was used to filter all data. For the Plk1 inhibitor/synchronized dataset, normalization was disabled. For the siRNA dataset, normalization was enabled. For both datasets imputation wasdisabled and single hit proteins were excluded. Un-paired student’s t-tests were conducted and corrected p-values (q-values) were calculated in Spectronaut. The mass spectrometry proteomics data have been deposited to the ProteomeXchange Consortium via the PRIDE partner repository with the dataset identifier PXD046039”.

### Molecular Biology

Plasmids for Cyclin F, WT PLK1, WT βTrCP1, and WT βTrCP2 were obtained from the human ORFeome v5.1 and cloned into the indicated pDEST vectors using gateway recombination. All AKAP2 vectors including N-Myc-AKAP2 WT (pcDNA 3.1+), N-Myc-AKAP2 DSG1 (pcDNA 3.1+), N-Myc-AKAP2 DSG2 (pcDNA 3.1+), N-Myc-AKAP2 DSG3 (pcDNA 3.1+), N-FLAG-AKAP2 WT (pGenLenti), N-FLAG-AKAP2 DSG2 (pGenLenti), AKAP2 WT (pGenDONR), and AKAP2 DSG2 (pGenDONR) were purchased from GenScript. An empty donor vector (attL1+2_pGenDONR) was also purchased from GenScript. AKAP2 WT and DSG2 pGenDONR plasmids were subcloned into pInducer20 using gateway recombination. An empty pInducer20 vector (EV) suitable for lentivirus production + transduction was generated using gateway recombination with the empty attL1+2_pGenDONR vector. AKAP2 WT pGenDONR plasmid was also cloned into the indicated pDEST vectors using gateway recombination. Site mutants including FLAG-PLK1 (T210A), FLAG-PLK1 (Pincer), FLAG-βTrCP1 (WD40^Mut^), and FLAG-βTrCP2 (WD40^Mut^) were generated using the Q5 site-directed mutagenesis kit (NEB; Cat. #E0552S). The βTrCP1 WD40^Mut^ changes amino acid 510 from R>A based on isoform 1 in uniprot (R474A when referring to isoform 2 in uniprot). Likewise, the βTrCP2 WD40^Mut^ changes amino acid 413 from R>A based on isoform 2 in uniprot (R447A when referring to isoform 1 in uniprot). Primers used for mutagenesis are described in **Table S5**.

### Lentiviral Particle Production in 293T Cells

To produce lentiviral particles for infection, HEK293T cells were co-transfected with a lentiviral plasmid (pGenLenti AKAP2 WT, pGenLenti AKAP2 DSG2, pInducer20 EV, pInducer20 AKAP2 WT, or pInducer20 AKAP2 DSG2) along with the lentiviral packaging plasmids VSV-G, Gag-pol, TAT, and Rev (packaging plasmids transfected at a 1:1:1:1 ratio). After 24 hours, cells were washed 1x with complete media and replaced with 10 mL fresh, complete media. After an additional 24 hours, the media was collected and centrifuged at 200×*g* for 3 minutes. Media was aliquoted and frozen at -80°C for at least 24 hours prior to transducing cells.

### Cell Line Generation

HCT116 sgCCNF1 and HCT116 sgCCNF2 cell lines were generated by transducing parental HCT116 cells (ATCC) with previously described CCNF lentiCRISPRv.2 constructs (Choudhury et al. 2016) and single cell populations, generated by serial dilution, were screen by immunoblot to confirm loss of cyclin F expression.

To produce HCT116 cells stably expressing FLAG-AKAP2 WT or FLAG-AKAP2 DSG2, cells were transduced with a 1:1 ratio of fresh complete media plus lentivirus-containing media with 8 µg/mL polybrene. 48 hours after transduction, cells were selected using 1 µg/mL puromycin for ∼7 days. Cell lines transduced with pInducer20 constructs were generated in the same manner as above, except selected for using 600 µg/mL G418 Sulfate for ∼7-10 days.

### Immunoprecipitation

For immunoprecipitation experiments, tagged protein constructs were expressed in HEK293T cells for 24 hours using transient transfection. After 24 hours, cells were washed once in 1x dPBS and harvested in PBS using a sterile cell lifter. Cells were then pelleted by centrifugation at 3,000×*g* for 3 minutes at 4C. Cell pellets were washed with 1x dPBS, lysed on ice for 15 minutes using NETN lysis buffer, and lysates were clarified by centrifugation at maximum speed (20,000×*g*) for 10 minutes at 4°C using a benchtop microcentrifuge. Protein concentration of samples was determined and normalized using Bradford Assay. Prior to IP, 10% of the total protein was removed as the input. The remaining lysate was then added to 30-50 µL of Anti-FLAG M2 (Sigma; cat. #F2426) affinity gel to isolate FLAG-tagged proteins (affinity gel was washed 3x using NETN lysis buffer prior to addition of cell lysate). Samples were immunoprecipitated for 2-4 Hrs at 4°C while rotating. After 2-4 Hrs, affinity gel was washed 3x at 4°C while rotating for 5 minutes each using NETN lysis buffer. After the final wash, affinity gel was suspended in 2x Laemmli sample buffer and boiled at 95°C for 10 minutes using a dry water-bath to elute proteins. Co-IP was assessed by immunoblotting, as described.

### AKAP2 IP-MS

For interactome analysis of AKAP2, Myc-EV or Myc-AKAP2 was transfected into HEK293T cells, in triplicate. Cells were collected and lysed in NETN lysis buffer, as described above. Lysates were snap frozen 2x using liquid nitrogen and then lysates were clarified by centrifugation at maximum speed (20,000×*g*) for 10 minutes at 4°C using a benchtop microcentrifuge. Protein concentration was determined and normalized using Bradford assay. Samples were immunoprecipitated using EZview c-Myc affinity gel (Sigma; cat. E6654). After IP, samples were washed 3x using NETN lysis buffer followed by 3 washes using 1x dPBS. Beads were covered in 1x dPBS and frozen at –80C until further analysis.

Immunoprecipitated protein samples were subjected to on-bead trypsin digestion as previously described (Rank et al. 2021). Briefly, after the last wash step of the immunoprecipitation, beads were resuspended in 50µl of 50mM ammonium bicarbonate, pH 8. On-bead digestion was performed by adding 1µg trypsin and incubated with shaking, overnight at 37°C. The following day, 1ug trypsin was added to each sample and incubated shaking, at 37°C for 3 hours. Beads were pelleted and supernatants were transferred to fresh tubes. The beads were washed twice with 100µl LC-MS grade water, and washes were added to the original supernatants. Samples were acidified by adding TFA to final concentration of 2%, to pH ∼2. Peptides were desalted using peptide desalting spin columns (Thermo), lyophilized, and stored at -80°C until further analysis.

Samples were analyzed by LC-MS/MS using an Easy nLC 1200 coupled to a QExactive HF mass spectrometer (Thermo). Samples were injected onto an Easy Spray PepMap C18 column (75 μm id × 25 cm, 2 μm particle size; Thermo) and separated over a 2 hour method. The gradient for separation consisted of 5–42% mobile phase B at a 250 nl/min flow rate, where mobile phase A was 0.1% formic acid in water and mobile phase B consisted of 0.1% formic acid in 80% ACN. The QExactive HF was operated in data-dependent mode where the 15 most intense precursors were selected for subsequent fragmentation. Resolution for the precursor scan (m/z 375–1700) was set to 60,000, while MS/MS scans resolution was set to 15,000. The normalized collision energy was set to 27% for HCD. Peptide match was set to preferred, and precursors with unknown charge or a charge state of 1 and ≥ 7 were excluded.

### Data analysis for AKAP2 IP-MS

Raw data files were searched against the reviewed human database (containing 20,396 entries), appended with a contaminants database, using Andromeda within MaxQuant. Enzyme specificity was set to trypsin, up to two missed cleavage sites were allowed, and methionine oxidation and N-terminus acetylation were set as variable modifications. A 1% FDR was used to filter all data. Match between runs was enabled (5 min match time window, 20 min alignment window), and a minimum of two unique peptides was required for label-free quantitation using the LFQ intensities.

Perseus was used for further processing (Tyanova et al. 2016). Only proteins with >1 unique+razor peptide were used for LFQ analysis. Proteins with 50% missing values were removed and missing values were imputed from normal distribution within Perseus. Log2 fold change (FC) ratios were calculated using the averaged Log2 LFQ intensities of sample compared to control, and students t-test performed for each pairwise comparison, with p-values calculated. Proteins with significant p-values (<0.05) and Log2 FC >1 were considered biological interactors.

### *In Vivo* Ubiquitination Assay

The *in vivo* ubiquitination assay was performed as described previously (Bonacci et al. 2014). Briefly, HEK293T cells were transfected with the indicated plasmids and harvested in PBS 48 hours post-transfection using a sterile cell lifter. Cells were treated with 20 µM MG132 and 20 µM PR-619 for the final 4 hours prior to harvesting. 20% of the cell suspension was removed to prepare inputs using the cell lysis procedure as described above. The remaining 80% of the cell suspension was lysed in 6M guanidine-HCl buffer and His_6_-tagged proteins were captured on Ni^2+^-NTA resin (Qiagen; cat. #30210). Pull-down eluates and inputs were separated on SDS-PAGE gels and analyzed by immunoblot, as described above.

### *In Vitro* Ubiquitination Assay

Recombinant UBA1, Ub, UBE2R2, UBE2L3, ARIH1, and CUL1/RBX1 were expressed and purified as previously described (Kamadurai et al. 2013; Scott et al. 2016).

Neddylation of CUL1/RBX1 (N8-CUL1/RBX1) was performed similar to (Duda et al. 2008). In short, all proteins were transformed into BL21-Codon Plus (DE3)-RIL *E. coli*, grown at 37°C, and expressed overnight after the addition of 0.6mM IPTG. Proteins were purified at 4°C via affinity chromatography, tags were removed via treatment with protease where applicable, and proteins were further purified by size-exclusion chromatography into assay buffer (20mM Hepes pH 8.0, 200mM NaCl, 1mM DTT). Proteins were flash-frozen in small aliquots using liquid nitrogen and stored at -80°C.

p4489 Flag-βTRCP was a gift from Peter Howley (Addgene plasmid # 10865) (Zhou et al. 2000). A βTRCP truncation (amino acids 175-C, hereafter referred to as βTRCP) was cloned into a modified pRSF vector that contains an N-terminal his-MBP tag. Untagged SKP1 was likewise cloned into pET3a using Gibson Assembly. BL21-Codon Plus (DE3)-RIL *E. coli* were co-transformed with both plasmids and grown in LB containing 150µg/ml ampicillin, 50µg/ml kanamycin, and 30µg/ml chloramphenicol. Protein expression was induced with 0.6mM IPTG at 18°C overnight. Cells were isolated by centrifugation, resuspended in 50mM HEPES pH 7.5, 200mM NaCl, 5mM β-ME, and 10mM imidazole, lysed by sonication, and clarified by centrifugation. Lysate was passed over Ni-NTA resin and the complex was eluted with lysis buffer supplemented with 250mM imidazole. βTRCP/SKP1 was dialyzed overnight, concentrated, and subjected to size-exclusion chromatography over an SD200 into assay buffer.

Fluorescent substrate peptides were synthesized by Biomatik based on previous studies to 95% purity (Scott et al. 2016). Peptides were dissolved to 1mM in 50mM HEPESpH 8.0, 50mM NaCl. Peptide sequences are as follows:

β-Catenin WT: Ac-KAAVSHWQQQSYLDSGIHSGATTAPRRASY-(PEG)2-K^TAMRA^

β-Catenin pSpS: Ac-KAAVSHWQQQSYLD(pS)GIH(pS)GATTAPRRASY-(PEG)2-K^TAMRA^

Akap2 WT: Ac-KDDDHGILDQFSRSVNVSLTQEELDSGLDELSVRS-(PEG)2-K^TAMRA^

Akap2 pS: Ac-KDDDHGILDQFSRSVNVSLTQEELD(pS)GLDELSVRS-(PEG)2-K^TAMRA^

For in vitro ubiquitination assays, proteins and peptides were diluted to 10x final concentration in assay buffer. Final reaction conditions are as follows: 0.1µM UBA1, 1µM UBE2R2, 0.75µM UBE2L3, 0.4µM MBP-βTRCP/SKP1, 0.5µM ARIH1, 0.5µM N8-CUL1/RBX1, 100µM Ub, and 0.25µM TAMRA-peptides. Reaction mixtures were moved to room temperature for 5 minutes before 5mM MgCl2 and 5mM ATP was added to start the reaction. Time points were quenched in 4x SDS-buffer, loaded onto 4-12% Bis-Tris SDS-PAGE gels, and scanned at the Cy3 channel on an Amersham Typhoon. Gel image is representative of 4 independent experiments.

### GO Analysis

All GO analysis was performed using Metascape (Zhou et al. 2019).

### Data Processing

40% contrast was added to all immunoblot images. Contrast was applied uniformly to all images and does not alter the interpretation of the data.

### BioRender Images

Illustrations in figures 1A, S5A, and 7A were created using BioRender.com

### Data Availability

Processed proteomics data files are available as supplemental tables.

All raw proteomics data files will be made publicly available on ProteomeXchange upon acceptance. During review, proteomics data can be accessed by reviewers and journal editors using the following information:

Project accession: PXD046039

Username: Password: 7bi00Bhp

## Acknowledgements

We thank lab members and colleagues at UNC for helpful discussions throughout this project. We thank Prasad Jallepalli (Memorial Sloan Kettering) for generously providing the RPE1-PLK1^WT^ and RPE1-PLK1^AS^ cell lines. The Emanuele lab (RM, TB, XW, CH and MJE) is supported by the UNC University Cancer Research Fund (UCRF), the National Institutes of Health (R01GM120309, R01GM134231), and the America Cancer Society (Research Scholar Grant; RSG-18-220-01-TBG). RM was also supported by NCI T32 CA071341. DLB was supported by NIH T32GM008570. TB was supported by F31CA189328. The Brown lab (DLB, KCR and NGB) is supported by NIH R35GM128855 and UCRF. This research is based in part upon work conducted using the UNC Proteomics Core Facility, which is supported in part by NCI Center Core Support Grant (2P30CA016086-45) to the UNC Lineberger Comprehensive Cancer Center. We thank Thomas Webb and Natalie Barker for their contributions to the proteomics work performed in this manuscript.

## Author Contributions

RM and ME conceived of experiments.

RM performed most cell and molecular experiments.

CH, XW, TB, EDT, and OGC helped with cell and molecular biology experiments.

DLB, NGB, and ME designed reagents for *in vitro* assays. DLB and KCR purified proteins and DLB performed assay under supervision from NGB and ME.

CAM and LEH assisted in designing proteomics experiments and performed and helped analyze data for all mass spectrometry experiments.

TBB generated HCT116 cyclin F KO cell lines.

RM and ME discussed, drafted, and wrote the manuscript.

All authors provided feedback on the final draft.

