## Supplemental Figs for "Proteomic Analysis Reveals a PLK1-Dependent G2/M Degradation Program and Links PKA-AKAP2 to Cell Cycle Control"

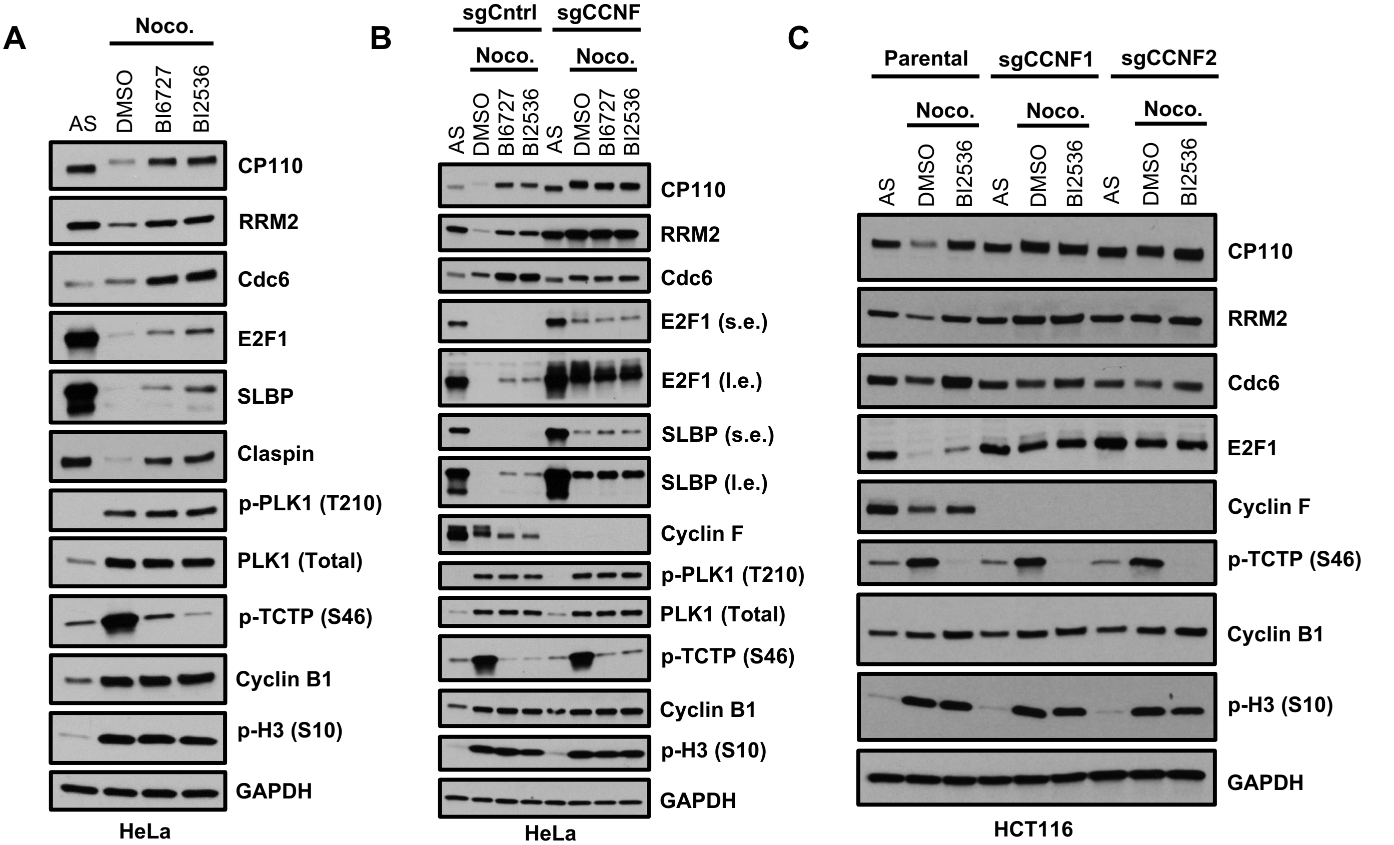


**Figure S1. PLK1 Regulates the Degradation of $SCF^{Cyclin\ F}$ Substrates**

(A) HeLa cells were grown asynchronously or were co-treated with 100 ng/mL nocodazole plus DMSO, 100 nM BI6727, or 100 nM BI2536 for 16 hours. After 16 hours, mitotic-arrested cells were collected using mitotic "shake-off" procedure. All cell lysates were analyzed by immunoblot for the indicated proteins.

(B) Experiment performed as in (A) except performed in either control HeLa cells or in HeLa cells where the Cyclin F gene has been deleted using CRISPR/Cas9.

(C) Experiment performed as in (A) except performed in either parental HCT116 cells or in HCT116 cells where the Cyclin F gene has been deleted using CRISPR/Cas9.

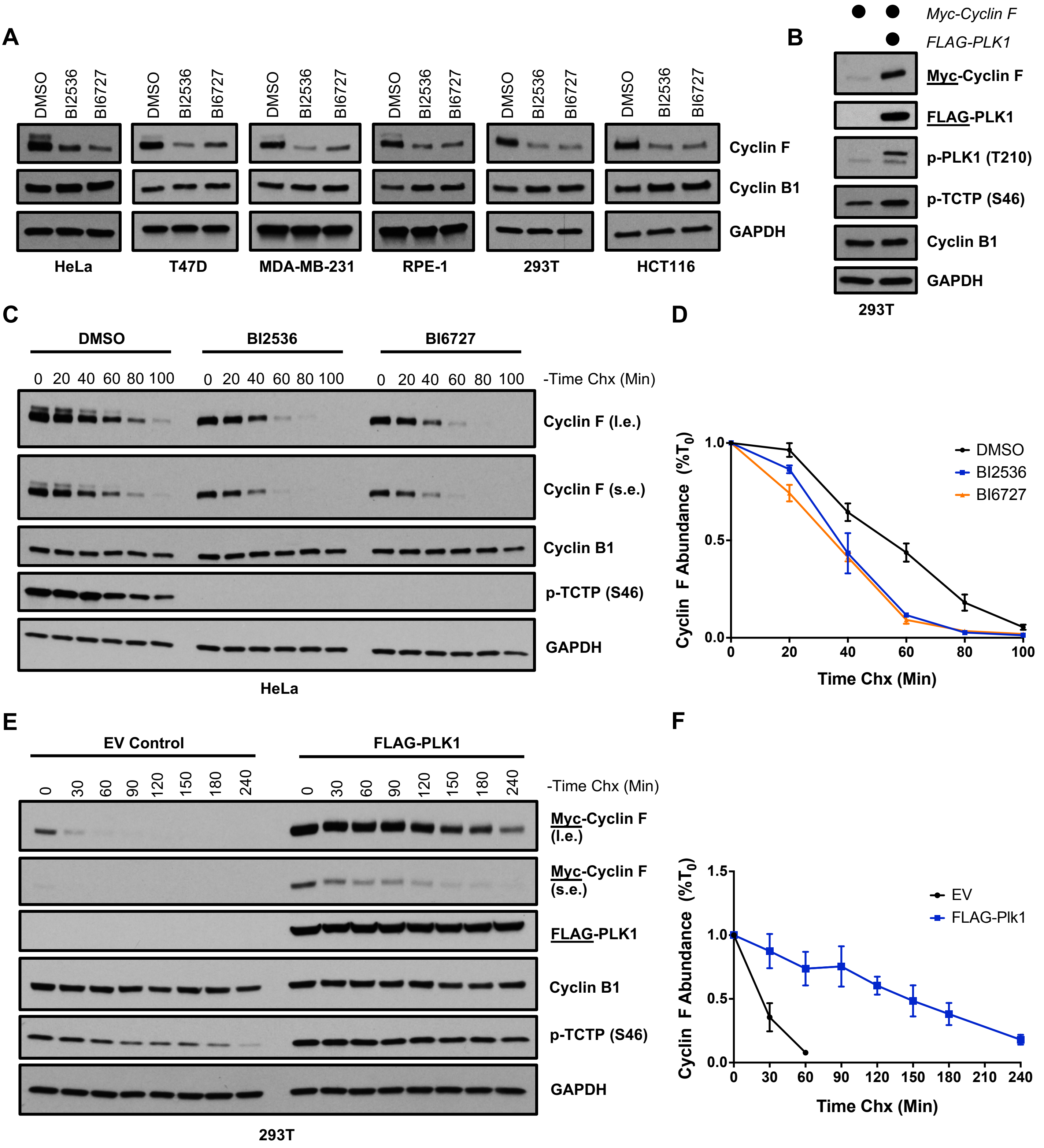


**Figure S2. PLK1 Regulates the Abundance and Stability of Cyclin F**

(A) The indicated cell lines were treated with DMSO or with 10 μM BI2536 or BI6727 for 4 hours. After 4 hours all cell lysates were analyzed by immunoblot for the indicated proteins.

(B) HEK293T cells were co-transfected with Myc-Cyclin F together with an empty vector (lane 1) or with FLAG-PLK1 (lane 2) for 24 hours. 24 hours post-transfection, cells were collected and all cell lysates were analyzed by immunoblot for the indicated proteins.

(C) HeLa cells were treated with DMSO, 10 μM BI2536, or 10 μM BI6727 for 1 hr. After 1 hour, cells were treated with 100 μg/mL of the protein synthesis inhibitor cycloheximide (Chx) in the continuous presence of the PLK1 inhibitors and cells were collected every 20 minutes for 100 minutes. All cell lysates were analyzed by immunoblot for the indicated proteins.

(D) Quantification of experiment performed in (C). Quantification was performed in FIJI and data shown are mean +/- SEM for n = 3 experiments.

(E) HEK293T cells were co-transfected with Myc-Cyclin F together with an empty vector or with FLAG-PLK1 for 24 hours. 24 hours post-transfection cells were treated with 100 μg/mL of the protein synthesis inhibitor cycloheximide (Chx) and cells were collected at various timepoints up to 240 minutes. All cell lysates were analyzed by immunoblot for the indicated proteins.

(F) Quantification of experiment performed in (E). Quantification was performed in FIJI and data shown are mean +/- SEM for n = 3 experiments.

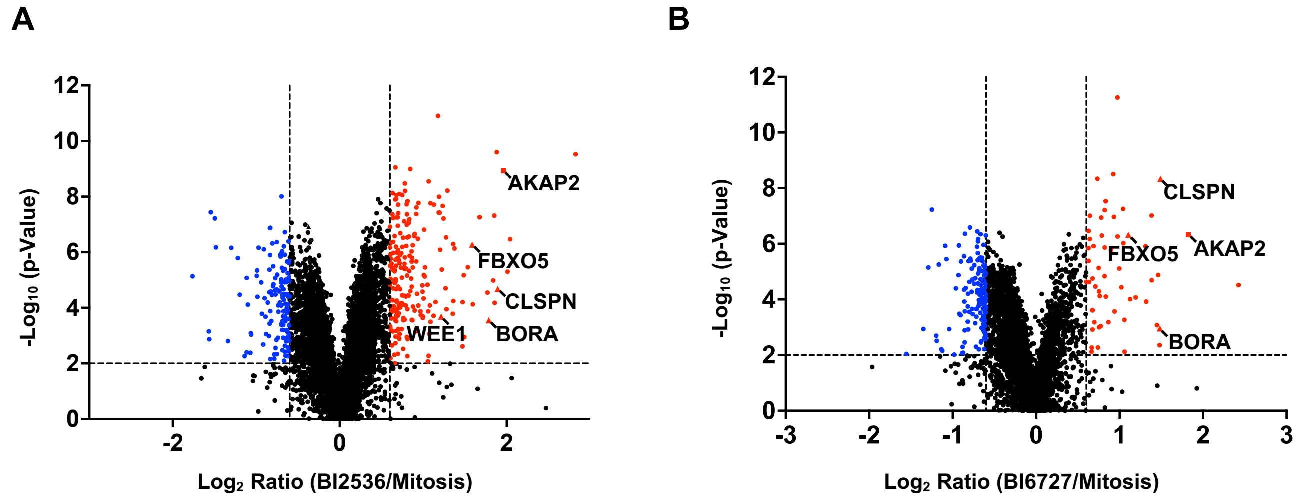


**Figure S3. Mass-Spectrometry Analysis shows that AKAP2 Expression is PLK1-Dependent during Mitosis**

(A) Same plot as shown in Figure 1D with AKAP2 indicated

(B) Volcano plot showing the $Log_2$ ratio (x-axis) and $-Log_{10}$ (p-value) (y-axis) of all proteins identified by mass-spectrometry when comparing samples from BI6727-treated mitotic cells to samples from DMSO-treated mitotic cells. Significant fold-change cutoffs were set at $Log_2 = -0.6$ or $Log_2 = 0.6$ and statistical significance was set at $-Log_{10}$ (p-value) = 2 (p = 0.01) which are denoted by the dashed lines. Positive controls are labeled and indicated as triangle dots.

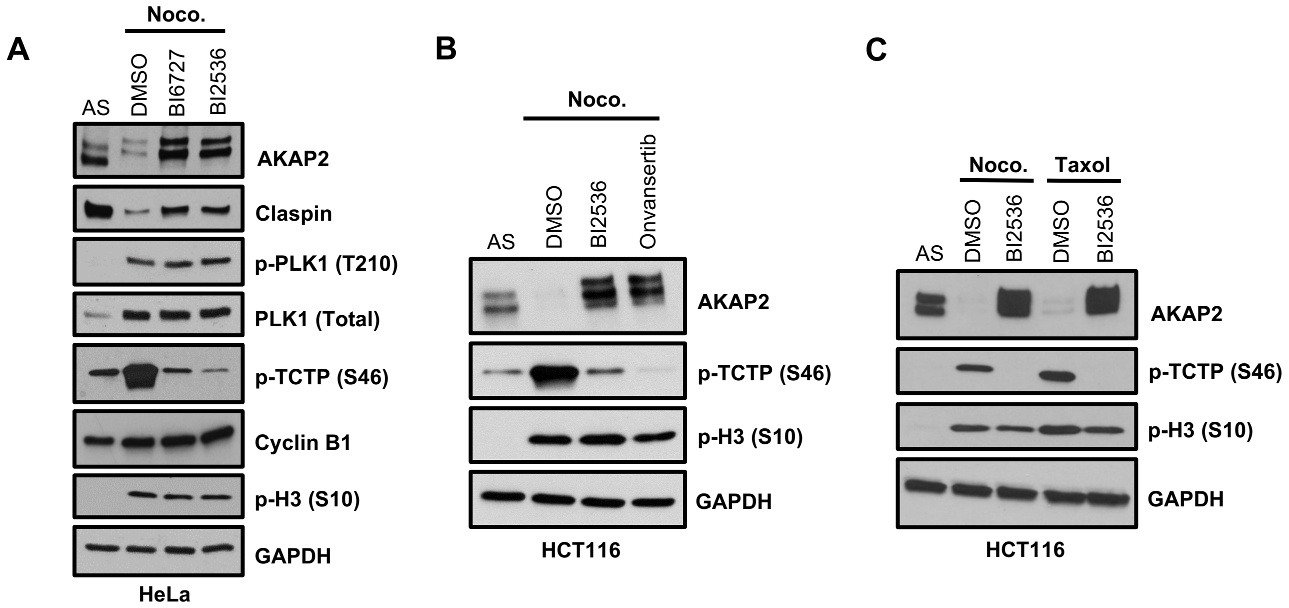


**Figure S4. Further Validation that AKAP2 abundance is reduced during Mitosis in a PLK1-dependent manner**

(A) HeLa cells were grown asynchronously or were co-treated with 100 ng/mL nocodazole plus DMSO, 100 nM BI6727, or 100 nM BI2536 for 16 hours. After 16 hours, mitotic-arrested cells were collected using mitotic “shake-off” procedure. All cell lysates were analyzed by immunoblot for the indicated proteins.

(B) HCT116 cells were grown asynchronously or were co-treated with 100 ng/mL nocodazole plus DMSO, 100 nM BI2536, or 250 nM Onvansertib for 16 hours. After 16 hours, mitotic-arrested cells were collected using mitotic “shake-off” procedure. All cell lysates were analyzed by immunoblot for the indicated proteins.

(C) HCT116 cells were grown asynchronously or were co-treated with 100 ng/mL nocodazole plus DMSO or 100 nM BI2536 or were co-treated with 50 nM Taxol plus DMSO or 100 nM BI2536 for 16 hours. After 16 hours, mitotic-arrested cells were collected using mitotic “shake-off” procedure. All cell lysates were analyzed by immunoblot for the indicated proteins.

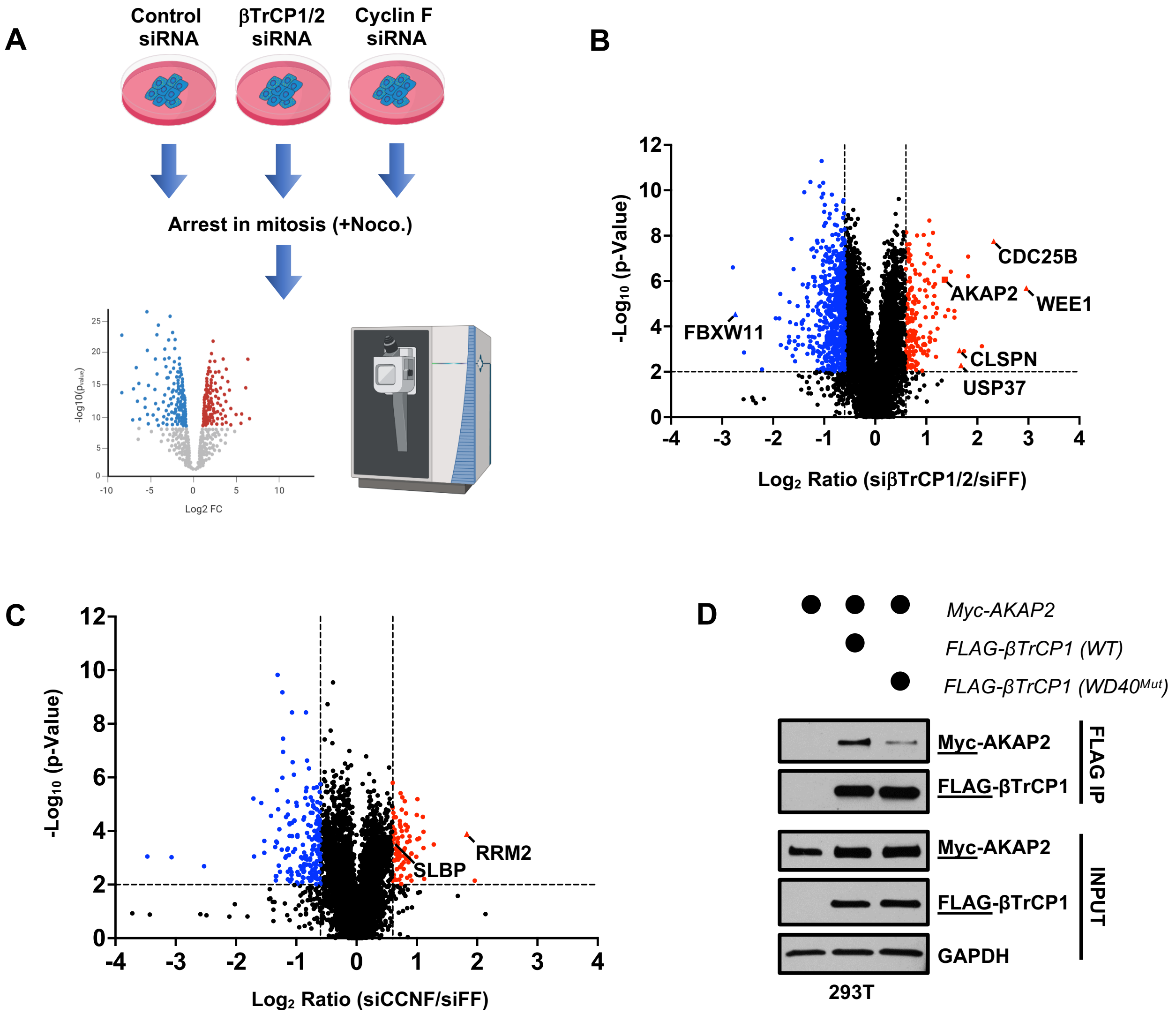


**Figure S5. βTrCP is the E3 Ubiquitin Ligase for AKAP2 During Mitosis**

(A) Schematic depicting the workflow for the siRNA proteomics experiment. HCT116 cells were transfected with siRNA targeting βTrCP1/2, CCNF, or firefly luciferase (siFF) as a non-targeting control for 48 hours. During the final 16 hours, cells were treated with 100 ng/mL nocodazole. After 16 hours, mitotic-arrested cells were collected using mitotic “shake-off” procedure. All samples were then prepared for label free, quantitative LC-MS/MS analysis using data independent acquisition (DIA).

(B) Volcano plot showing the $Log_2$ ratio (x-axis) and $-Log_{10}$ (p-value) (y-axis) of all proteins identified by mass-spectrometry when comparing samples from siβTrCP-treated mitotic cells to samples from siFF-treated mitotic cells. Significant fold-change cutoffs were set at $Log_2 = -0.6$ or $Log_2 = 0.6$ and statistical significance was set at $-Log_{10}$ (p-value) = 2 (p = 0.01) which are denoted by the dashed lines. Positive controls are labeled and indicated as triangle dots.

(C) Volcano plot showing the $Log_2$ ratio (x-axis) and $-Log_{10}$ (p-value) (y-axis) of all proteins identified by mass-spectrometry when comparing samples from siCCNF-treated mitotic cells to samples from siFF-treated mitotic cells. Significant fold-change cutoffs were set at $Log_2 = -0.6$ or $Log_2 = 0.6$ and statistical significance was set at $-Log_{10}$ (p-value) = 2 (p = 0.01) which are denoted by the dashed lines. Positive controls are labeled and indicated as triangle dots.

(D) HEK293T cells were co-transfected with plasmids expressing Myc-AKAP2 together with an empty vector control (lane 1), wild-type FLAG-βTrCP1 (lane 2), or a version of FLAG-βTrCP1 with a mutation in the substrate binding domain ($WD40^{Mut}$; lane 3). 24 hours post-transfection, cells were collected and βTrCP was immunoprecipitated using anti-FLAG affinity gel. Eluates were analyzed by immunoblot for the indicated proteins.

**A**

**AKAP2 'DSG1'**

| | |
|---|---|
| AKAP2 (*Homo sapiens*) | 379-PPE**DSG**AS**A**AKG-390 |
| AKAP2 (*Mus musculus*) | 400-TLE**DGG**TQ**A**AKE-411 |
| PALM2-AKAP2 (*Xenopus laevis*) | 588-QSI**QGE**TP**L**VKA-599 |
| AKAP2 'DSG1'Mutant | 379-PPE**AAA**AS**A**AKG-390 |

**B**

**AKAP2 'DSG3'**

| | |
|---|---|
| AKAP2 (*Homo sapiens*) | 499-NIS**DSG**AS**N**ETT-510 |
| AKAP2 (*Mus musculus*) | 520-NIS**DSG**AS**N**ETP-431 |
| PALM2-AKAP2 (*Xenopus laevis*) | 699-NIS**DSG**AP**N**ETM-710 |
| AKAP2 'DSG3'Mutant | 499-NIS**AAA**AS**N**ETT-510 |

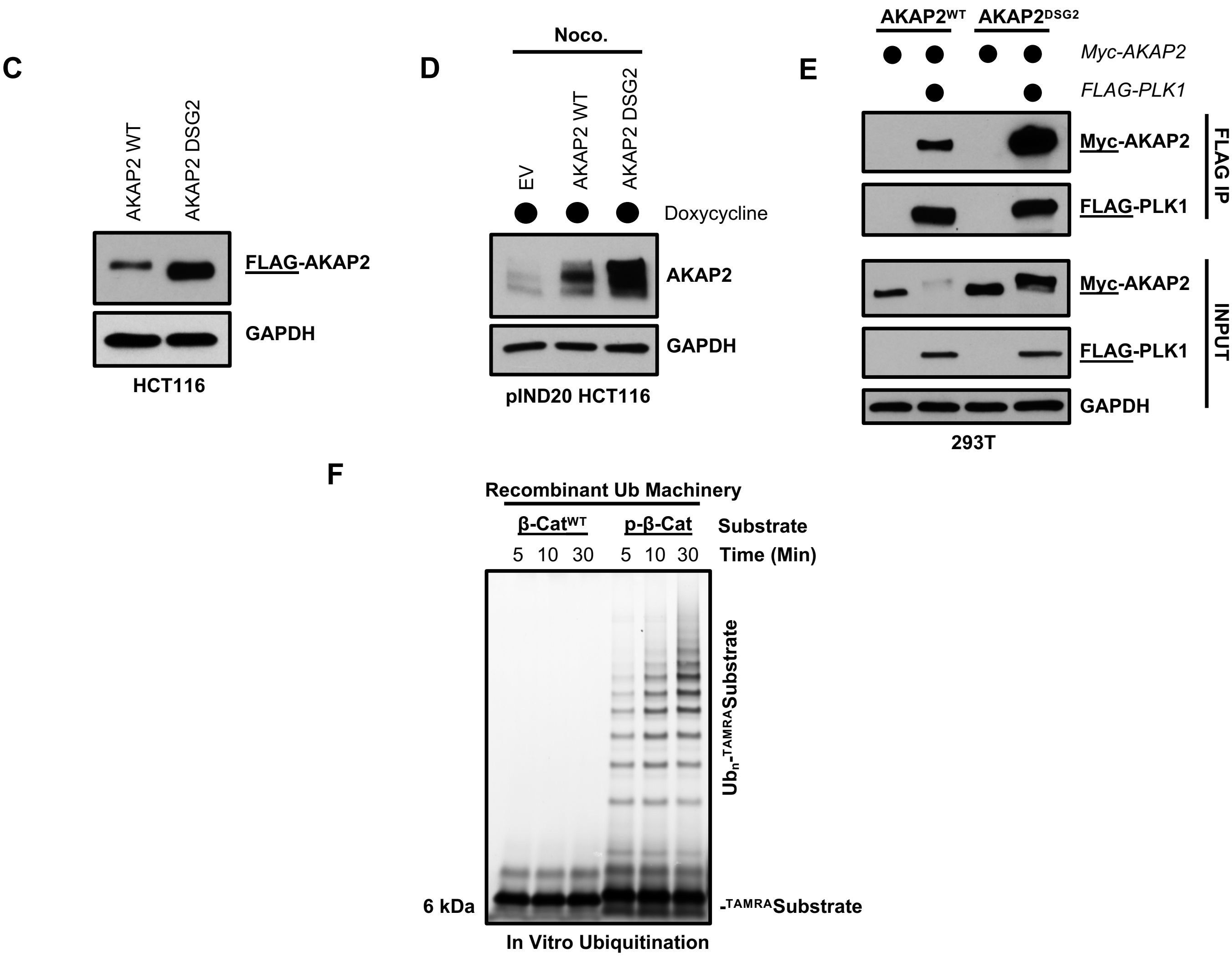


**Figure S6. Identification of Functional βTrCP Degron in AKAP2**

(A) Sequence alignment of the 'DSG1' motif in *Homo sapiens, Mus musculus,* and *Xenopus laevis.* The sequence of the 'DSG1' motif AKAP2 mutant used in future experiments is shown below.

(B) Sequence alignment of the 'DSG3' motif in *Homo sapiens, Mus musculus,* and *Xenopus laevis.* The sequence of the 'DSG3' motif AKAP2 mutant used in future experiments is shown below.

(C) Asynchronously growing HCT116 cells stably expressing FLAG-AKAP2$^{WT}$ or FLAG-AKAP2$^{DSG2}$ were collected and cell lysates were analyzed by immunoblot for the indicated proteins.

(D) HCT116 cells were engineered to express empty-vector, AKAP2$^{WT}$, or AKAP2$^{DSG2}$ under the control of a TET-inducible promoter. Cells were co-treated with 25 ng/mL doxycycline and 100 ng/mL nocodazole for 18 hours. After 18 hours, mitotic-arrested cells were collected using mitotic "shake-off" procedure. All cell lysates were analyzed by immunoblot for the indicated proteins.

(E) HEK293T cells were co-transfected with plasmids expressing Myc-AKAP2 (WT or "DSG2' degron mutant) together with an empty vector control or FLAG- PLK1. 24 hours post-transfection, cells were collected and PLK1 was immunoprecipitated using anti-FLAG affinity gel. Eluates were analyzed by immunoblot for the indicated proteins.

(F) Control *in vitro* ubiquitination reaction using TAMRA-labeled β-Catenin WT or phospho-peptide as a substrate, monitored by fluorescent scanning of an SDS-PAGE gel. Representative of n=4 independent experiments. Experiment is from the same gel as Figure 5G.

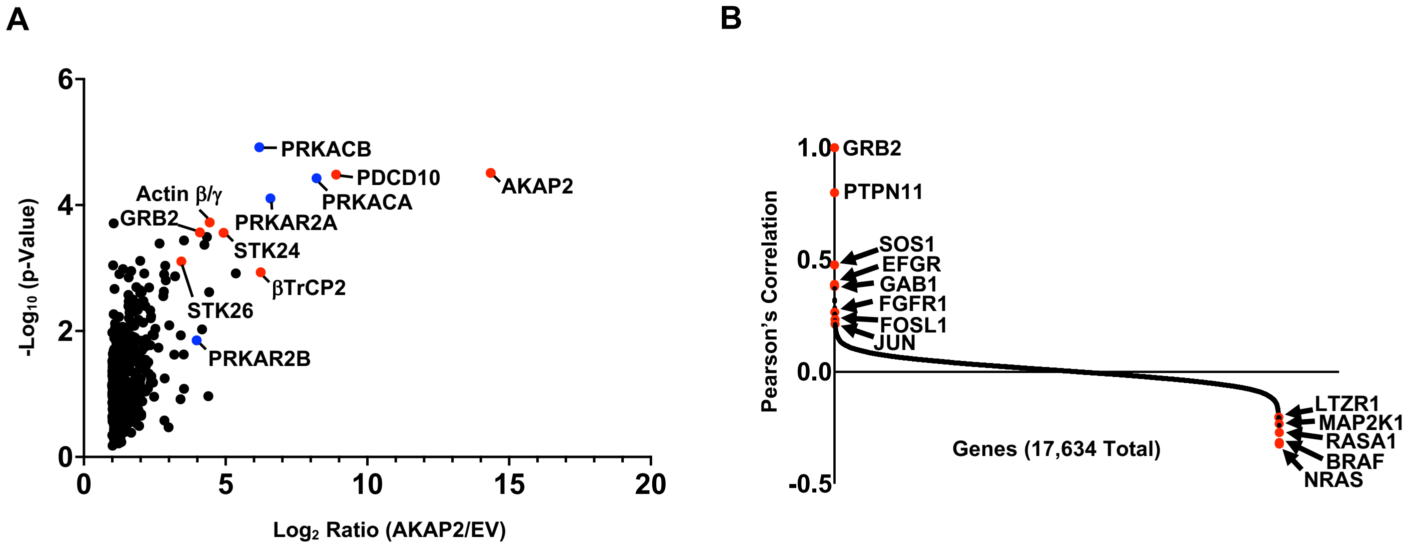


**Figure S7. IP-MS Analysis Reveals the AKAP2 Interactome**

(A) Plot showing the $Log_2$ ratio (x-axis) and $-Log_{10}$ (p-value) (y-axis) of all proteins identified by mass-spectrometry when comparing samples from AKAP2 IP to samples from EV control IP. PKA subunits are known interactors of AKAP proteins and are indicated in blue.

(B) DepMap analysis of GRB2 showing that it is highly correlated with genes that function within the MAPK and AKT signaling pathways.
